# The Acute Myeloid Leukemia variant DNMT3A Arg882His is a DNMT3B-like enzyme

**DOI:** 10.1101/720557

**Authors:** Allison B. Norvil, Lama AlAbdi, Bigang Liu, Nicole E. Forstoffer, Amie R. Michie, Taiping Chen, Humaira Gowher

## Abstract

Mutations in DNMT3A, particularly the Arg882His substitution is highly prevalent in acute myeloid leukemia. Although the reduced activity of DNMT3A Arg882His variant alters DNA methylation, the underlying cause of its oncogenic effect is not fully understood. Our data show that DNMT3A Arg882His variant acquires CpG flanking sequence preference highly similar to that of DNMT3B. Interestingly, a similar substrate preference was observed in DNMT3A WT enzyme upon the loss of cooperative kinetic mechanism. We tested if DNMT3A Arg882His could preferably methylate DNMT3B-specific target sites. Rescue experiments in *Dnmt3a/3b* double knockout mouse embryonic stem cells show that the corresponding Arg878His mutation in mouse DNMT3A severely impairs its ability to methylate major satellite DNA, a DNMT3A-preferred target, but has no overt effect on the ability to methylate minor satellite DNA, a DNMT3B-preferred target. Our data suggest that methylation of DNMT3B target sites by DNMT3A Arg882His variant could contribute to its oncogenic potential.

## Introduction

DNA methylation in mammals is critical for development and maintenance of the somatic cell state^1,2^. It has diverse functions including regulation of gene expression and silencing of repetitive elements^3–5^. In mammals, CpG methylation is established and maintained by DNA methyltransferases (DNMTs)^6^. DNMT1 largely functions as a maintenance methyltransferase by copying the methylation pattern from parent to daughter strand during DNA replication^7,8^. The DNMT3 family includes two active homologs DNMT3A and DNMT3B, and an inactive homolog DNMT3L^9,10^. These enzymes perform *de novo* DNA methylation, which predominantly takes place during early embryogenesis and pluripotent stem cell differentiation^9,11,12^. Whereas DNMT3A is expressed ubiquitously, DNMT3B is highly expressed during early embryogenesis and largely silenced in somatic cell types^13–15^. Despite having a high sequence similarity (>40%), DNMT3A and DNMT3B have distinct preferences for some target sites, although many sites can be methylated redundantly^16^. This is represented by a preference of DNMT3A for major satellite repeats, whereas DNMT3B preferentially targets minor satellite repeats^9,11^.

Mutations in DNMT3A and DNMT3B have been identified in several diseases. Germline transmitted mutations of DNMT3A and DNMT3B cause Tatton-Brown-Rahman syndrome and ICF (Immunodeficiency, Centromeric instability, and Facial anomalies) syndrome, respectively^9,17–20^. Somatic mutations in DNMT3A are commonly found in patients with acute myeloid leukemia (AML) and other hematologic neoplasms^21,22^. Detailed studies revealed that ∼20% of AML cases have heterozygous DNMT3A mutations, with the majority (∼60-70%) carrying the Arg882His mutation, which cause genome-wide hypomethylation^21,23–25^. Given that genetic knockout of one copy of DNMT3A exhibits no obvious phenotype, the heterozygous DNMT3A Arg882His mutation was suggested to have a dominant negative effect^9,21^. The observation that the expression of the murine DNMT3A Arg878His variant (equivalent to Arg882His in human DNMT3A) in mouse embryonic stem cells (mESCs) causes genome-wide loss of DNA methylation further supported the dominant negative activity of this variant^26,27^. In contrast, DNMT3B mutations have not been identified in cancer. However aberrant overexpression of DNMT3B is highly tumorigenic, including in AML^14,28–30^.

Overexpression of DNMT3B in AML leads to disease prognosis similar to that in patients with DNMT3A Arg882His mutation^29,30^.

The effect of Arg882His in the activity of DNMT3A could be predicted from the crystal structure of the DNMT3A catalytic domain (DNMT3A-C) showing that it forms homodimers and tetramers through two interaction surfaces^31,32^. Several AML-associated DNMT3A mutations, including Arg882His, which are present at or close to the protein-protein interface disrupt tetramer formation and lead to reduced catalytic activity *in vitro*^21,27,33–36^. Further, DNMT3A-C tetramers can oligomerize on DNA, forming nucleoprotein filaments^37,38^. This oligomerization allows the enzyme to bind and methylate multiple CpG sites in a cooperative manner, thus increasing its activity^39^. Our previous biochemical studies show that in DNMT3A-C, the Arg882His substitution results in loss of cooperativity potentially causing a decrease in its catalytic activity^40^. Interestingly, in DNMT3B, which is a non-cooperative enzyme, the homologous mutation, Arg829His, has no effect on its catalytic activity *in vitro*^40^. A recent co-crystal structure of the DNMT3A-C/DNA complex shows that Arg882 interacts with the phosphate backbone of the DNA at the N+3 position downstream of the CpG site^41^. The DNMT3A-C Arg882His variant was shown to have a bias for G at N+3 when compared to the wild-type (WT) enzyme^42^.

In this study, we tested whether this flanking sequence preference was directly due to the substitution of Arg with His or caused by loss of cooperative mechanism in the variant enzyme. Our data show that all variants of Arg882 found in AML patients have low catalytic activity and lack a cooperative mechanism. We further show that in the absence of cooperative mechanism at low enzyme concentration, the DNMT3A-C WT enzyme prefers G at the N+3 position, similar to that of DNMT3A-C Arg882His variant. Based on previous observations that DNMT3B-C is a non-cooperative enzyme, we performed a comparative analysis of its substrate preference with that of the DNMT3A-C WT and the Arg882His variant. DNA methylation levels at 56 CpGs were rank ordered to compute consensus motifs that are preferred by these enzymes. Interestingly, our data show that the Arg882His variant and DNMT3B-C have a similar preference for nucleotides at N+1, 2 and 3 positions flanking the CpG site. These data strongly support that the gain of flanking sequence preference is due to loss of cooperative mechanism, and suggest that the DNMT3A-C Arg882His variant could methylate DNMT3B-C preferred sites. We tested this “gain of function” prediction by expressing WT mouse DNMT3A or the Arg878His (corresponding to Arg882His in human DNMT3A) in *Dnmt3a/3b* double knockout (DKO) mESCs. Our data show that whereas the DNMT3A Arg878His variant failed to rescue methylation at the major satellite repeats (DNMT3A preferred target sites), its ability to methylate the minor satellite repeats (DNMT3B preferred target sites), was comparable to that of DNMT3A WT enzyme. Further, our analysis of the CpG flanking sequences of the satellite repeats show that whereas major satellite repeats have an A or T at the N+3 position, the minor satellite repeats are enriched in G at N+3 position. This observation provides the previously unknown explanation for the substrate specificity of DNMT3A and DNMT3B for major and minor satellite repeats, respectively. Taken together, our data provide novel mechanistic insights into DNMT3A and DNMT3B substrate specificities that could influence the oncogenic potential of these enzymes. We suggest that in leukemia, DNMT3A Arg882His substitution establishes a DNMT3B-like activity, the tumorigenic properties of which are exploited by cancer cells.

## Results

### The cooperative kinetic mechanism is absent in DNMT3A Arg882 variants

The catalytic properties of the DNMT3A-C Arg882His variant have been previously characterized *in vitro* showing that the mutation reduces the activity and attenuates tetramerization of the enzyme^21,27,33^. DNMT3A-C WT methylates neighboring CpGs on DNA by a cooperative kinetic mechanism, in which the DNA bound tetramer promotes successive binding events through protein-protein interaction^39^. Our previous studies also show that the DNMT3A-C Arg882His variant fails to methylate DNA by a cooperative mechanism, which could be due to impaired tetramerization^26,33,40^. It is not clear, however, whether this is due to the loss of Arg or gain of His at this position. Based on the observation that many AML patients have mutations that lead to substitution of Arg882 to Cys or Ser, we asked if these variant enzymes could methylate multiple CpGs on a single DNA molecule in a cooperative manner. His-tagged recombinant DNMT3A-C (catalytic domain) WT and Arg882 variants were produced in *E. coli* and purified using Ni-NTA affinity chromatography to about 90 – 95% purity (Supplementary Fig. 1a)^43^. The catalytic activity of DNMT3A-C Arg882 variants was compared to the WT enzyme by performing DNA methylation assays using a 30-bp substrate containing one CpG site. Consistent with previous reports, a 60 - 80% loss of catalytic activity was observed in all variants compared to WT enzyme (Fig. 1a)^27,33,35,36,40,42^. Cooperativity is observed as an exponential relationship between catalytic activity enzyme concentrations shown for DNMT3A-C WT. We tested the cooperative mechanism of the Arg882 variants using the pUC19 plasmid. All Arg882 variant enzymes failed to methylate the substrate in a cooperative fashion (Fig. 1b). From these data, we conclude that the Arg882 residue plays a key role in the cooperative mechanism of DNMT3A-C.

**FIGURE 1.**
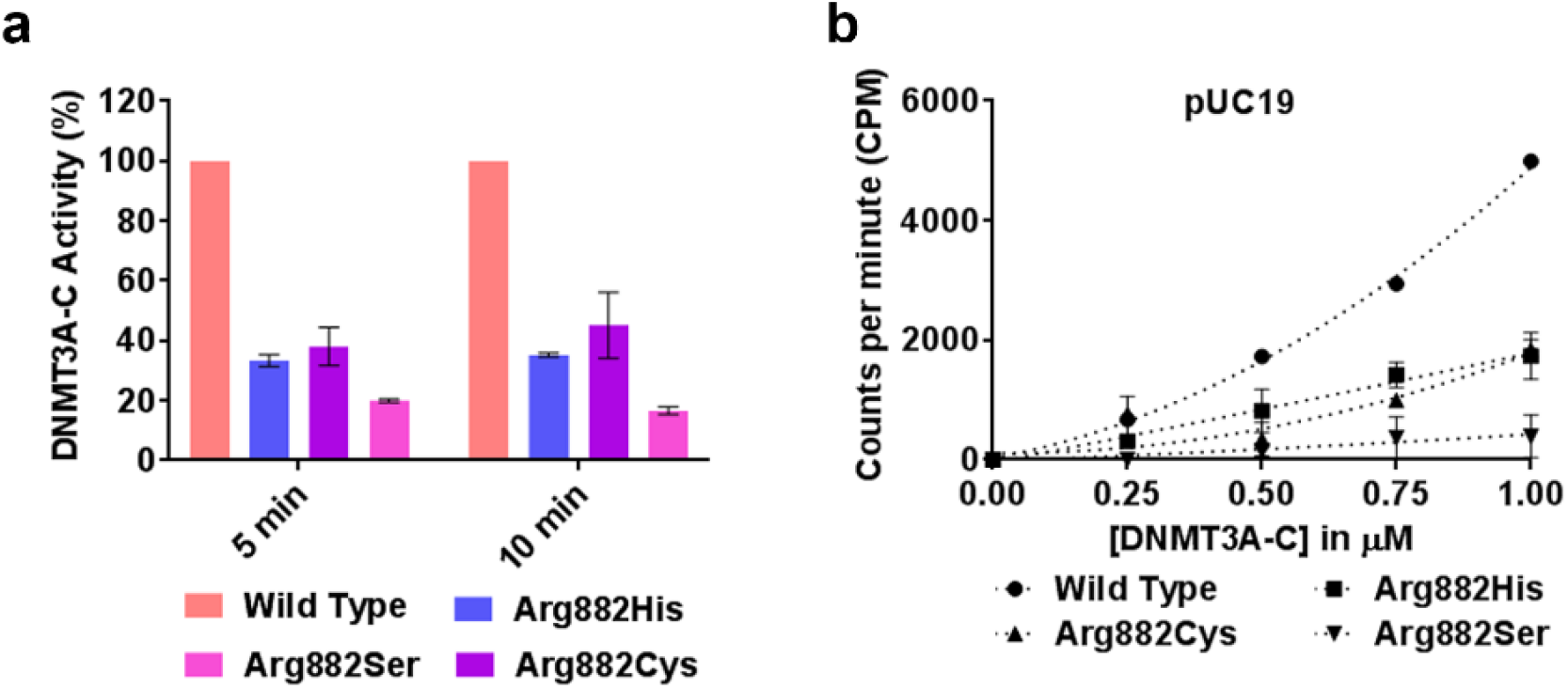
Relative activity and kinetic mechanism of DNMT3A-C WT and Arg882 variants. **a** Methylation activity of 1 μM DNMT3A-C WT and Arg882 variants was measured for 5 and 10 minutes using ^3^[H] labelled S-Adenosylmethionine (AdoMet). The transfer of radiolabeled –CH_3_ group to DNA was measured as counts per minute (CPM) using the MicroBeta scintillation counter. **b** Methylation activity of DNMT3A-C WT and Arg882 variants was measured for 10 minutes using 100 ngs of pUC19 plasmid as a substrate, at concentrations of enzymes varying from 0.25 to 1 μM. The enzymes were pre-incubated with DNA for 10 minutes at room temperature and the reaction was initiated by addition of AdoMet. Each data point is an average and standard error of the mean (*n*≥3 independent experiments).The data shows reduced activity and loss of cooperativity for all the variant enzymes.

### Loss of cooperativity modulates flanking sequence preference of DNMT3A

A recent study of the crystal structure of DNMT3A shows that the Arg882 residue interacts with the phosphate backbone of the nucleotide at N+3 position (N=CpG) and contributes to DNA binding^41^. It was shown recently that the Arg882His has a preference for G at the N+3 position compared to the WT enzyme^42^. Given that the Arg882 residue is also necessary for the cooperative kinetic mechanism of DNMT3A, we tested the relationship between loss of cooperative mechanism and flanking sequence preference. We performed *in vitro* methylation of a 509-bp DNA substrate containing 56 CpG sites using the WT and Arg882His variant enzymes (Supplementary Fig. 2). DNA methylation was quantified by bisulfite conversion and high throughput sequencing. Our data confirmed the previously reported G preference at N+3 for Arg882His variant compared to the WT enzyme (Fig. 2a). We have previously shown that at a low concentration (0.25 μM), DNMT3A-C does not multimerize on the 509-bp DNA substrate and therefore cannot methylate multiple CpGs using cooperative mechanism, which is observed at a higher concentration (1 μM) of the enzyme^40^. DNA methylation assays were performed using 0.25 μM and 1 μM enzyme, and the flanking sequence preferences were compared. The preference of sites by DNMT3A-C at 0.25 μM over 1 μM is represented by fold change greater than 1 (Fig. 2b, Supplementary Table 1). Strikingly, 3 out of 4 sites preferred by DNMT3A-C at 0.25 μM have G at N+3 position, which is similar to the flanking sequence preference of the DNMT3A-C Arg882His variant. When compared to the relative preference at the N+1 and N+2 positions, interestingly, the similarity was specifically observed at N+3 position, suggesting the role of cooperativity in modulating the interaction of the Arg882 residue with DNA.

**FIGURE 2.**
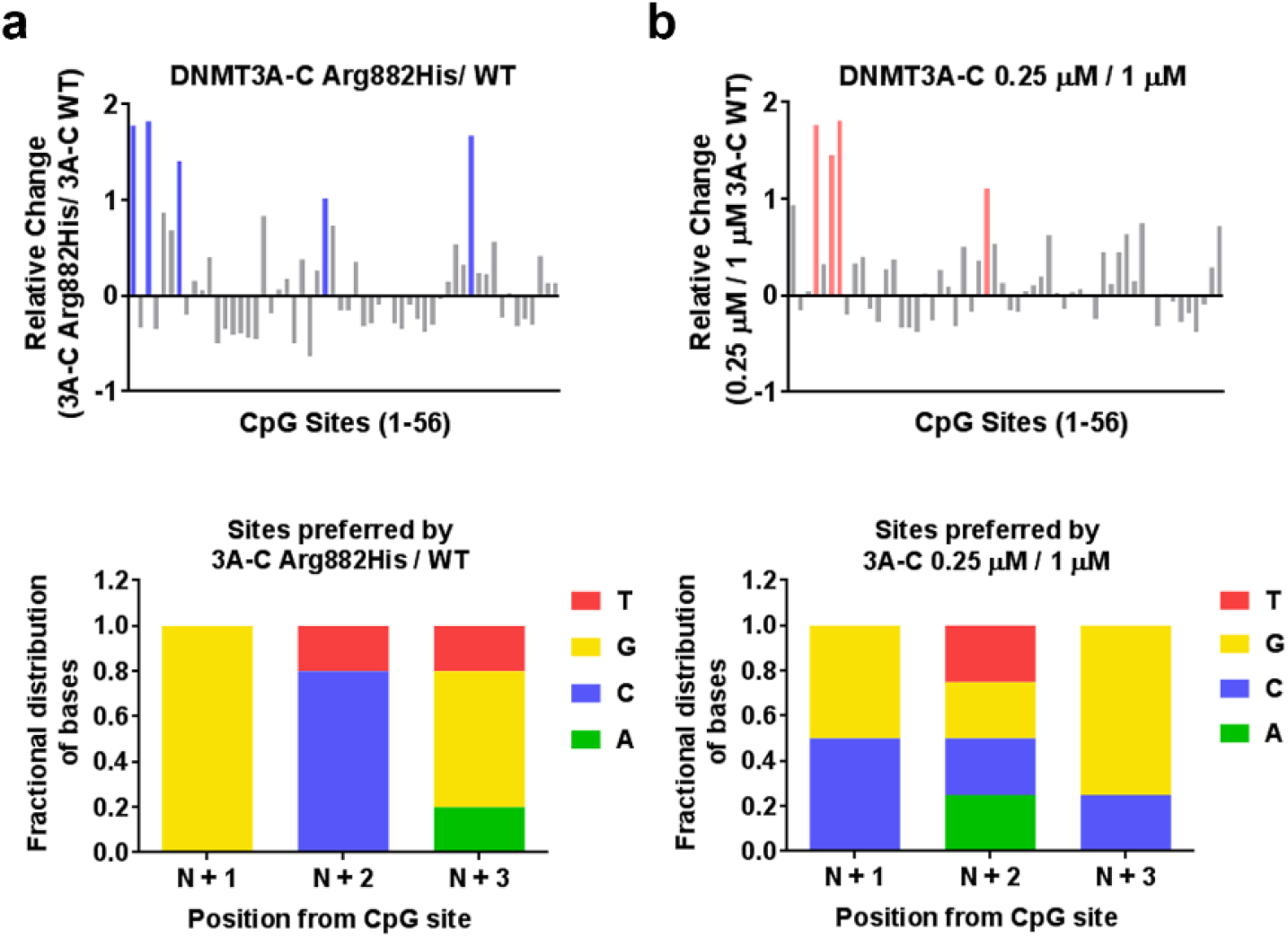
Effect of cooperativity on flanking sequence preference by DNMT3A-C. **a, b** DNA methylation of the 56 CpG sites in the 509-bp substrate was analyzed using bisulfite sequencing. Methylation reaction was carried out using 1 μM DNMT3A-C WT, and 1 μM DNMT3A-C Arg882His in **a**, and 0.25 μM and 1 μM DNMT3A-C WT in **b**. *Top panel* shows relative preference calculated for each CpG site (1-56) by the DNMT3A-C Arg882His compared to DNMT3A-C WT in **a**, and 0.25 μM compared to 1 μM of DNMT3A-C WT in **b**. Relative preference of 1 is equal to a 2-fold change, and is represented by blue and light pink bars, respectively. *Bottom panel* shows the fractional distribution of nucleotides at the preferred sites. Each bar represents nucleotides at positions N+1/2/3 respectively from the CpG site. The data show DNMT3A-C WT at low concentrations have flanking sequence preference similar to DNMT3A-C Arg882His variant.

### DNMT3B has a flanking sequence preference similar to DNMT3A Arg882His

DNMT3B is a homolog of DNMT3A that is frequently overexpressed in tumors, including AML^14,29^. Similar to DNMT3A-C Arg882His variant, DNMT3B-C functions as a non-cooperative enzyme^40^. Therefore, we asked if DNMT3B-C and the DNMT3A-C Arg882His variant have a similar flanking sequence preference. Although the nucleotide preference of DNMT3B is reported for N+1 position, the extended flanking sequence preference has not been thoroughly evaluated^44^. Using recombinant DNMT3B-C WT and the Arg829His variant (homologous to DNMT3A Arg882His), we performed *in vitro* methylation assays using the 509-bp DNA substrate. We compared the preferred sites of DNMT3B-C WT with DNMT3A-C WT, DNMT3B-C WT with DNMT3A-C Arg882His, and DNMT3B-C Arg829His with WT. The preferred sites of DNMT3B-C WT compared to DNMT3A-C WT showed pronounced increase at 11 sites, of which 8 had G at N+3 position (Fig. 3a). Strikingly, many of these sites also overlapped with those preferred by the DNMT3A-C Arg882His enzyme. This was confirmed by a direct comparison of the preferred sites of DNMT3B-C WT and the DNMT3A Arg882His enzyme, which showed only 4 sites preferred by DNMT3B-C over DNMT3A-C Arg882His (Fig. 3b). Interestingly, 3 out of the 4 sites carry G at N+3 position. The data also showed only a few sites that were preferred by DNMT3B-C the Arg829His variant compared to WT, suggesting that the Arg829 mutation has little or no effect on DNMT3B-C activity and flanking sequence preference (Fig. 3c). Unlike DNMT3A, DNMT3B methylates DNA using a processive kinetic mechanism^40^. Our data confirmed that, similar to DNMT3A-C WT, the Arg882His variant is a non-processive enzyme indicating that the Arg882His mutation only affects its flanking sequence preference (Supplementary Fig. 3). Taken together, these data suggest that the Arg882His substitution alters the specificity of DNMT3A to be similar to that of DNMT3B.

**FIGURE 3.**
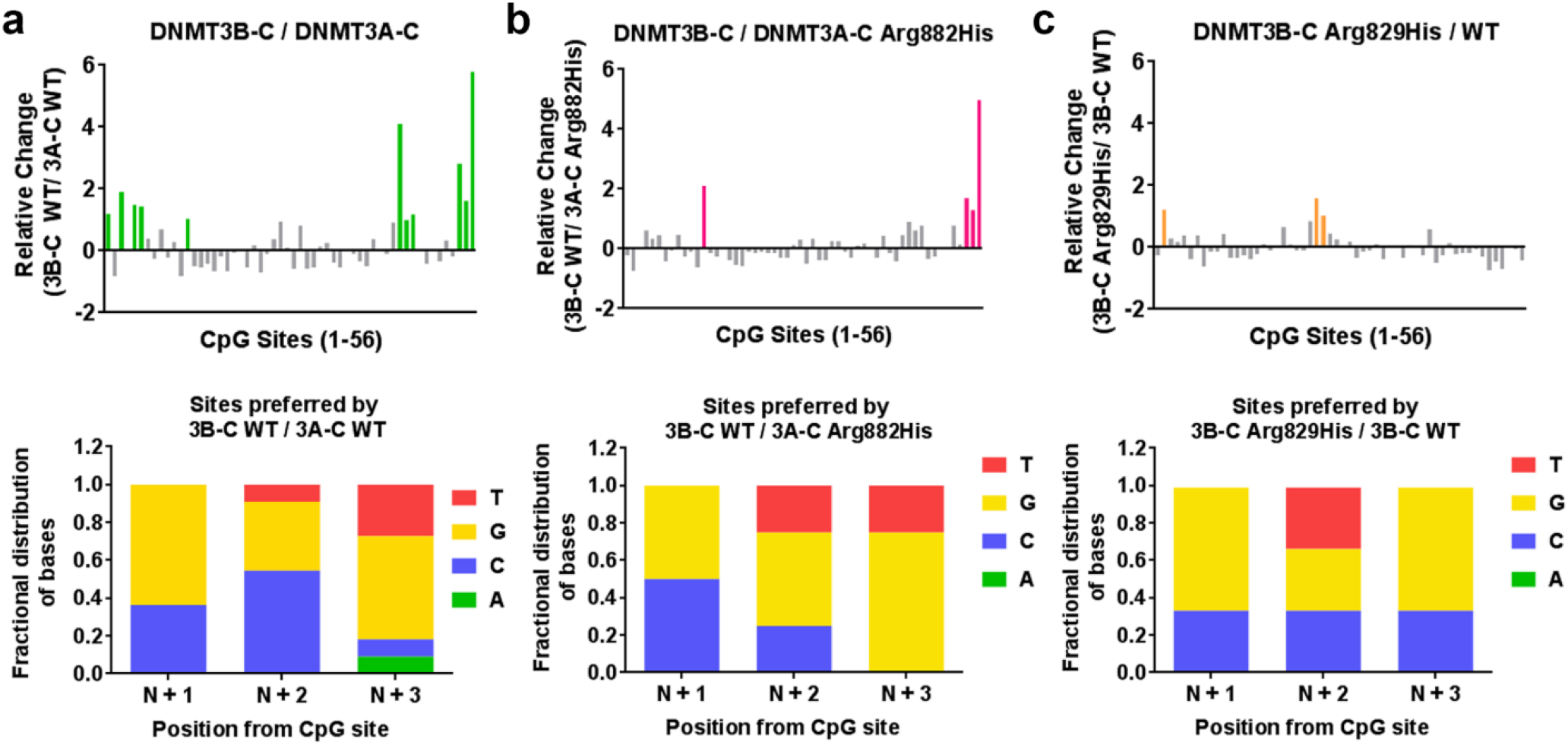
Comparative analysis of the flanking sequence preferences of DNMT3A-C Arg882His and DNMT3B-C WT**. a-c** DNA methylation of the 56 CpG sites in the 509-bp substrate was analyzed using bisulfite sequencing. Methylation reaction was carried out using 1 μM enzyme for 10 minutes. *Top panels* show relative preference calculated for each CpG site (1-56) by DNMT3B-C WT compared to DNMT3A-C WT in **a**, DNMT3B-C WT compared to DNMT3A-C Arg882His in **b**, and DNMT3B-C Arg829His compared to DNMT3B-C WT in **c**. Relative preference of 1 is equal to a 2-fold change, and is represented by green, light pink and orange bars, respectively. *Bottom panels* show the distribution of nucleotides at positions N+1/2/3 respectively from the CpG site at 11 preferred sites by DNMT3B-C WT in **a**, 4 preferred sites by DNMT3B-C WT in **b**, or the 3 sites preferred by DNMT3B-C Arg829His in **c**. The data show that substrate preference of DNMT3A-C Arg882His variant is similar to that of DNMT3B enzyme.

### The DNMT3A Arg882His variant acquires DNMT3B-like substrate preference

The methylation assays to determine the flanking sequence preference described above were performed for 10 minutes. which represents initial enzyme kinetics. To evaluate the substrate preference during multiple turnovers, the methylation assays were carried out for 30 and 60 minutes, allowing the enzyme kinetics to enter the steady state. Comparing site preference for DNMT3A-C Arg882His to DNMT3A-C WT, the data show progressive loss of preference for G at N+3 position from 30 to 60 minutes (Fig. 4a, b). This is expected under conditions where the enzyme, after methylating the preferred sites during the initial reaction, methylates other sites under multiple turnover conditions. A similar comparison of site preference of DNMT3B-C WT over DNMT3A-C WT or Arg882His variant, however, shows that the preference for G at N+3 position is maintained during 30- and 60-minute time points (Fig. 4b, c, e, f). This indicates that whereas DNMT3B-C has a strong intrinsic specificity for G at the N+3 position and methylates other sites at very low frequencies, the preference for this site is acquired by the DNMT3A-C Arg882His variant enzyme.

**FIGURE 4.**
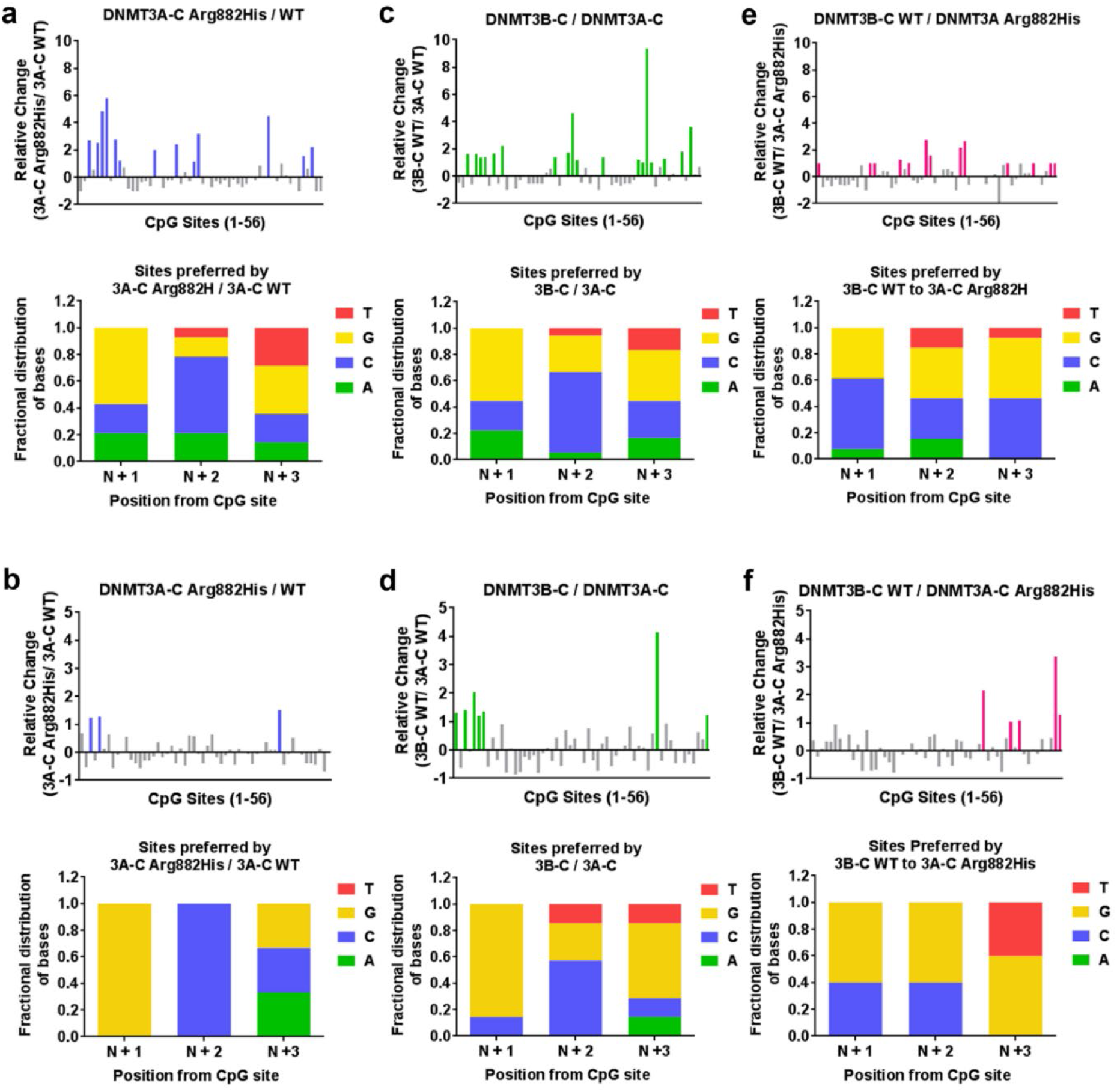
Flanking sequence preference at steady state kinetics. DNA methylation of the 56 CpG sites in the 509-bp substrate was analyzed using bisulfite sequencing. Methylation reactions were carried out using 1 μM enzyme for 30 minutes in **a, c, e**, and 60 minutes in **b, d, f**. *Top panels* show the relative preference calculated for each CpG site (1-56) by DNMT3A-C WT compared to the DNMT3A-C Arg882His variant in **a, b**, DNMT3B-C WT compared to DNMT3A-C WT in **c, d**, and DNMT3B-C WT compared to DNMT3A-C Arg882His in **e, f.** Relative preference of 1 is equal to a 2-fold change, and is represented by blue, green and pink bars, respectively. *Bottom panels* show the fractional distribution of nucleotides at the preferred sites. Each bar represents nucleotides at positions N+1/2/3 respectively from the CpG site. The data show that the similarity in flanking sequence preference between DNMT3A-C Arg882His and DNMT3B-C is maintained during steady state kinetics.

### Temporal change in flanking sequence preference by WT and variant DNMT3 enzymes

We extended the analysis of the flanking sequence preference to determine the preferred trinucleotides at the N+1/2/3 and N−1/2/3 positions by DNMT3A-C, DNMT3B-C and the variant enzymes. Based on the occurrence of different sites, the methylation at a site was calculated as a ratio of observed to expected fractional methylation. Comparing the flanking sequence preference of DNMT3A-C WT, DNMT3A-C Arg882His, DNMT3B-C WT, and DNMT3B-C Arg829His, the data showed that whereas DNMT3A-C Arg882His, DNMT3B-C WT, and DNMT3B-C Arg829His cluster together at 21 out of the 64-nucleotide combinations, DNMT3A-C WT shows an opposite and a distinct preference (Fig. 5a, Supplementary Fig. 5a, Supplementary Table 2).

**FIGURE 5.**
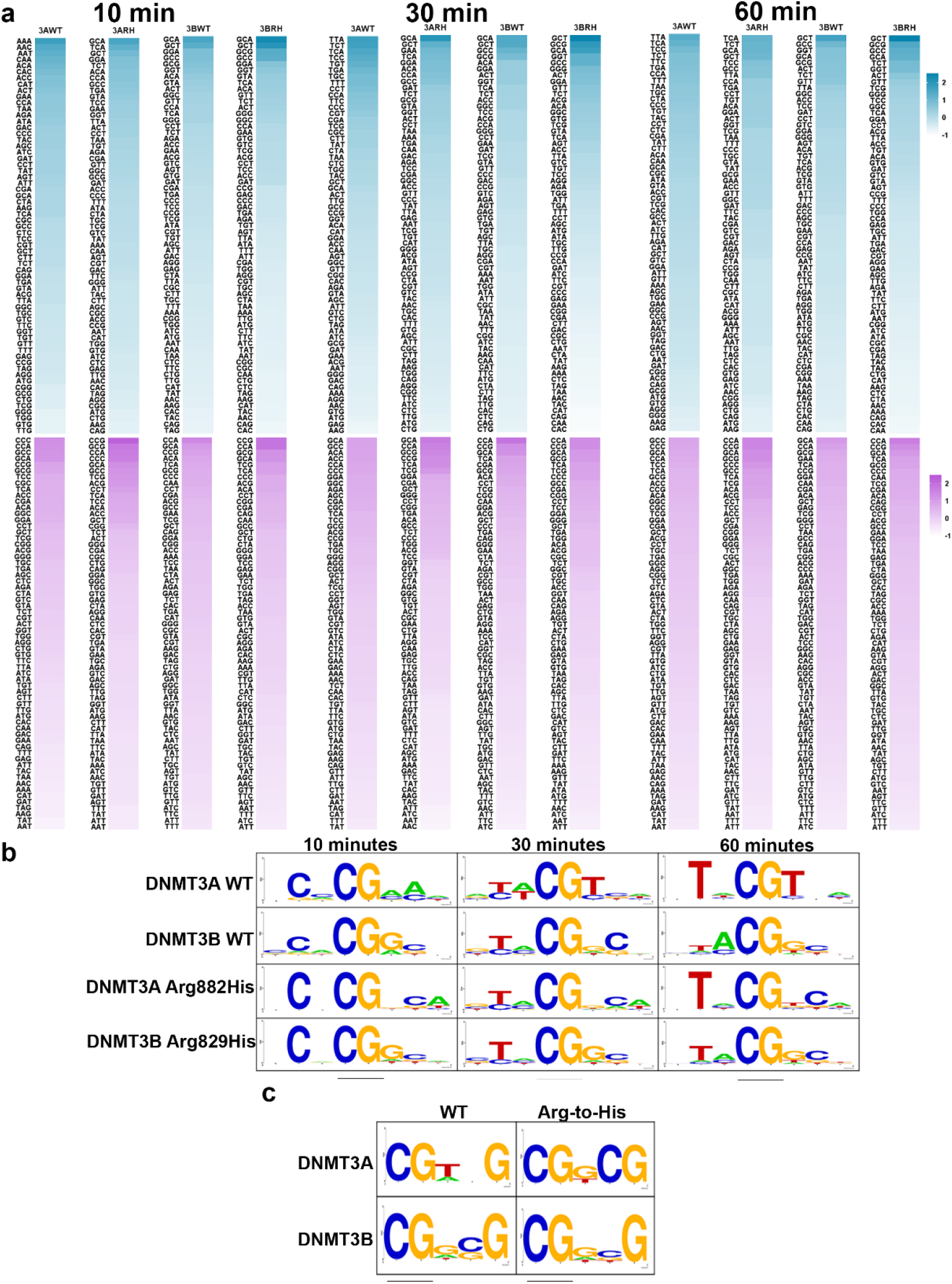
Trinucleotide sequence flanking CpG preferred by WT and variant DNMT3 enzymes. **a** Heat-map showing the preference of different trinucleotide sets by DNMT3A-C WT (3AWT), DNMT3A-C Arg882His (3ARH), DNMT3B-C WT (3BWT), or DNMT3B-C Arg829His (3BRH) at 10, 30, and 60 minutes. Sequences with values greater or equal to 1 are considered as preferred. Upper panels in blue represent the flanking sequence preference at the N +1/2/3 positions. Lower panels in purple represent the flanking sequence preference at the N-1/2/3 positions. **b** Consensus sequence generated by WebLogo from top ten preferred sequences by each enzyme as methylation reaction proceeds from 10 to 60 minutes. **c** Consensus flanking sequence for methylated sites with a G at the N+3 position. The data show that for the DNMT3A-C Arg882His variant the nucleotide preference at N+1 is lost and gained at N+3, while at the N+2 position it switches from an A to a C, making it very similar to the flanking sequence preference of DNMT3B-C. Similarly, the methylated sites with G at the N+3 position have inner flanking sequence similar between DNMT3A-C Arg882His and DNMT3B-C, whereas it is different for DNMT3A-C WT enzyme

Based on the top 10 preferred sites, we used WebLogo application^45^ to determine the consensus sequence logo flanking the CpG sites for each enzyme. The analysis was performed for methylation data collected at 10, 30, and 60 min time points to monitor the temporal order of site preference by these enzymes (Fig. 5b). As expected, the data again show a dramatic difference in flanking sequence preference between DNMT3A-C and DNMT3B-C. Whereas DNMT3A-C WT shows a preference for A or T at the N+1/2/3 positions, DNMT3B-C prefers G and C. The preferred sequence of the DNMT3A-C Arg882His variant is strikingly similar to that of DNMT3B-C, particularly at N+1, where it loses preference for T, and at N+2, where it gains a strong preference for C. The sites with A at the N+1 position are most preferred by DNMT3A-C WT and are methylated within the first 10 minutes, whereas sites with T at the N+1 position are methylated at 30 and 60 minutes (Fig. 5b, Supplementary Fig. 5a). This is consistent with previous studies which show DNMT3A to prefer T at N +1 from CpG site^44^. DNMT3A-C WT enzyme has least preference for sites with G at N+1 position, which is just the opposite for DNMT3B-C and DNMT3A-C Arg882His (Supplementary Fig. 5b, c). Both DNMT3B-C and DNMT3A-C Arg882His enzymes strongly prefer sites with C at N+2 positon and methylate sites with A and T at N+1 position slowly and with a weaker preference. Comparing preference of all four enzymes for nucleotides at position N-1/3 shows a weak or no preference, whereas at N-2, all of them show a strong preference for C and T (Fig. 5b). This is in agreement with crystal structure data showing fewer interactions between DNMT3A and the nucleotides upstream of CpG site^41^. Taken together, these data show that DNMT3A-C Arg882His has a similar nucleotide preference as DNMT3B-C WT, not only at N+3 but also at positions N+1 and N+2 from the CpG site.

We next selected CpG sites that had G at the N+3 position and generated the preferred flanking sequence logo using the WebLogo application^45^. A comparison between the consensus sequences again show a striking similarity between DNMT3A-C Arg882His, DNMT3B-C, and DNMT3B-C Arg829His, with G at N+1 position, which is the most preferred nucleotide. In contrast, G at N+1 is least preferred by DNMT3A-C WT enzyme. The consensus flanking sequence of DNMT3A-C WT had a T at this site. This uncovered the importance of the nucleotide at the N+1 position, which can affect the interaction of DNMT3 enzymes with DNA (Fig. 5c). We therefore tested if T at the N+1 position affects the preference for G at the N+3 position of DNMT3B-C and DNMT3A-C Arg882His enzymes, by using 30-bp oligonucleotide substrates which contain a central CpG site and varying combinations of nucleotides at N+2/3 or N-2/3 positions. The positions N+1 and N-1 were held constant with T and A respectively (Supplementary Table 3). Methylation assays using radiolabeled AdoMet were performed for 10 minutes and total methylation was measured. Our data show that the preference of DNMT3B-C and the DNMT3A-C Arg882 variant for G at N+3 position is lost (Supplementary Fig. 4a-d). DNMT3A-C and its variants show rather strong preference for sites with A at the N+2 position, whereas DNMT3B-C shows a weak preference for sites with A or C occupying the N+2 position (Supplementary Fig. 4e). Interestingly, DNMT3B-C Arg829His showed reduced activity when compared to the WT enzyme, indicating an adverse effect of T at N+1 position on its activity (Supplementary Fig. 4f). These data confirm our previous observations that the interaction of DNMT3A-C Arg882His at the N+3 position is strongly influenced by the nucleotide at the N+1 position.

### DNMT3A-C Arg882His and DNMT3B-C preferably methylate the same CpG site in the *Meis1* enhancer

We next tested if the change in flanking sequence preference of DNMT3A-C Arg882His affected methylation of the regions that are known to be spuriously hypomethylated in AML patients^46^. The *Meis1* gene is expressed during development and regulates leukemogenesis and hematopoiesis by promoting self-renewal of progenitor-like cells^47,48^. The enhancer of *Meis1* is methylated by DNMT3A during normal hematopoietic stem cell (HSC) differentiation, whereas in AML patients expressing the DNMT3A Arg882His variant, this region is largely hypomethylated^46,49^. A 1-kb region of the *Meis* enhancer was used as a substrate for methylation reactions, and methylation was analyzed by bisulfite sequencing (Supplementary Fig. 6a). The average methylation at all CpG sites showed an expected high level of methylation by DNMT3A-C WT compared to DNMT3A-C Arg882His and DNMT3B-C WT, and Arg882 variants showed a loss of cooperativity on this substrate *in vitro* (Supplementary Fig. 6b, c). Next, we computed the flanking sequence preference of these enzymes as described above. The data showed a strong preference of DNMT3B-C for one site compared to DNMT3A-C, which had G at N+1 position (Fig. 6a, Supplementary Fig. 6a). Interestingly, this site was also the most preferred site by DNMT3A-C Arg882His compared to DNMT3A-C WT (Fig. 6b). A comparison between DNMT3B-C and DNMT3A-C Arg882His confirmed that DNMT3B-C has a higher preference for this site (Fig. 6c). Although there are 3 sites in this substrate with G at N+3 position, these sites have either T, C, or A at N+1 position, which are weakly preferred by DNMT3B-C and the DNMT3A-C Arg882His variant. These data again confirm that DNMT3A-C Arg882His has acquired a substrate preference that is similar to that of DNMT3B-C. The data also show that the nucleotide at the N+1 position has substantial effect on the flanking sequence preference of DNMT3 enzymes.

**FIGURE 6.**
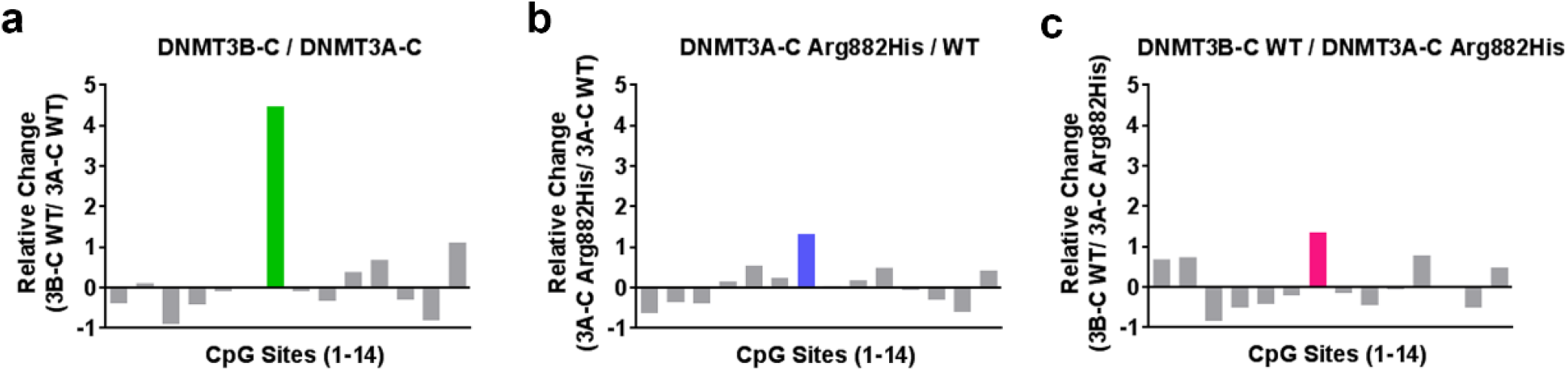
Relative activity and site preference of DNMT3 WT and mutant enzymes on *Meis1* enhancer substrate. A 1-kb *Meis1* enhancer region was used as substrate for *in vitro* methylation reactions by DNMT3A-C WT, the Arg882His variant, and DNMT3B-C WT. DNA methylation of the 14 CpG sites was analyzed using bisulfite sequencing. Relative preference for each of the 14 CpG sites was calculated for DNMT3B-C compared to DNMT3A-C in **a**, DNMT3A-C Arg882His compared to DNMT3A-C WT in **b**, and DNMT3B-C WT compared to DNMT3A-C Arg882His in **c**. Relative preference of 1 is equal to a 2-fold change, and is represented by green, blue and pink bars, respectively.. DNMT3A-C Arg882His and DNMT3B-C prefer to methylate same site in this substrate, however DNMT3B-C shows a very strong preference compared to the DNMT3A-C Arg882His variant.

### Mouse DNMT3A Arg878His retains its activity for minor satellite DNA in mESCs

While DNMT3A and DNMT3B redundantly methylate many genomic regions in cells, they also have preferred and specific targets^9,11,50^. For example, in murine cells, DNMT3A preferentially methylates the major satellite repeats in pericentric regions, whereas DNMT3B preferentially methylates the minor satellite repeats in centromeric regions^9,11^. Based on our observations, we predicted that DNMT3A Arg882His would prefer DNMT3B-specific targets. To test the idea, we carried out rescue experiments in late-passage *Dnmt3a/3b* DKO mESCs, which show severe loss of global DNA methylation, including at the major and minor satellite repeats^11^. mESCs express two major DNMT3A isoforms, DNMT3A1 (full length) and DNMT3A2 (a shorter form that lacks the N terminus of DNMT3A1), with both showing identical activity^11,26,51^. We transfected *Dnmt3a/3b* DKO mESCs with plasmid vectors and generated stable lines expressing mouse DNMT3A1 WT, DNMT3A1 Arg878His, DNMT3B1 WT, or DNMT3B1 Arg829His (catalytically inactive DNMT3A1 and DNMT3B1, with their PC motif in the catalytic center being mutated^52^, were included as negative controls) (Fig. 7a). The genomic DNA from these cell lines was harvested, and DNA methylation at the major and minor satellite repeats was analyzed by digestion with methylation-sensitive restriction enzymes followed by Southern blot. Consistent with our previous results^26^, the ability of DNMT3A1 to rescue methylation at the major satellite repeats is severely impaired with the Arg878His substitution (Fig. 7b). However, at the minor satellite repeats, which are largely methylated by DNMT3B, DNMT3A Arg878His rescues DNA methylation to levels similar to the DNMT3A WT enzyme (Fig. 7c). These data demonstrate that the DNMT3A AML mutant specifically retains its ability to methylate DNMT3B preferred target sites, while losing preference for sites methylated by DNMT3A.

**FIGURE 7.**
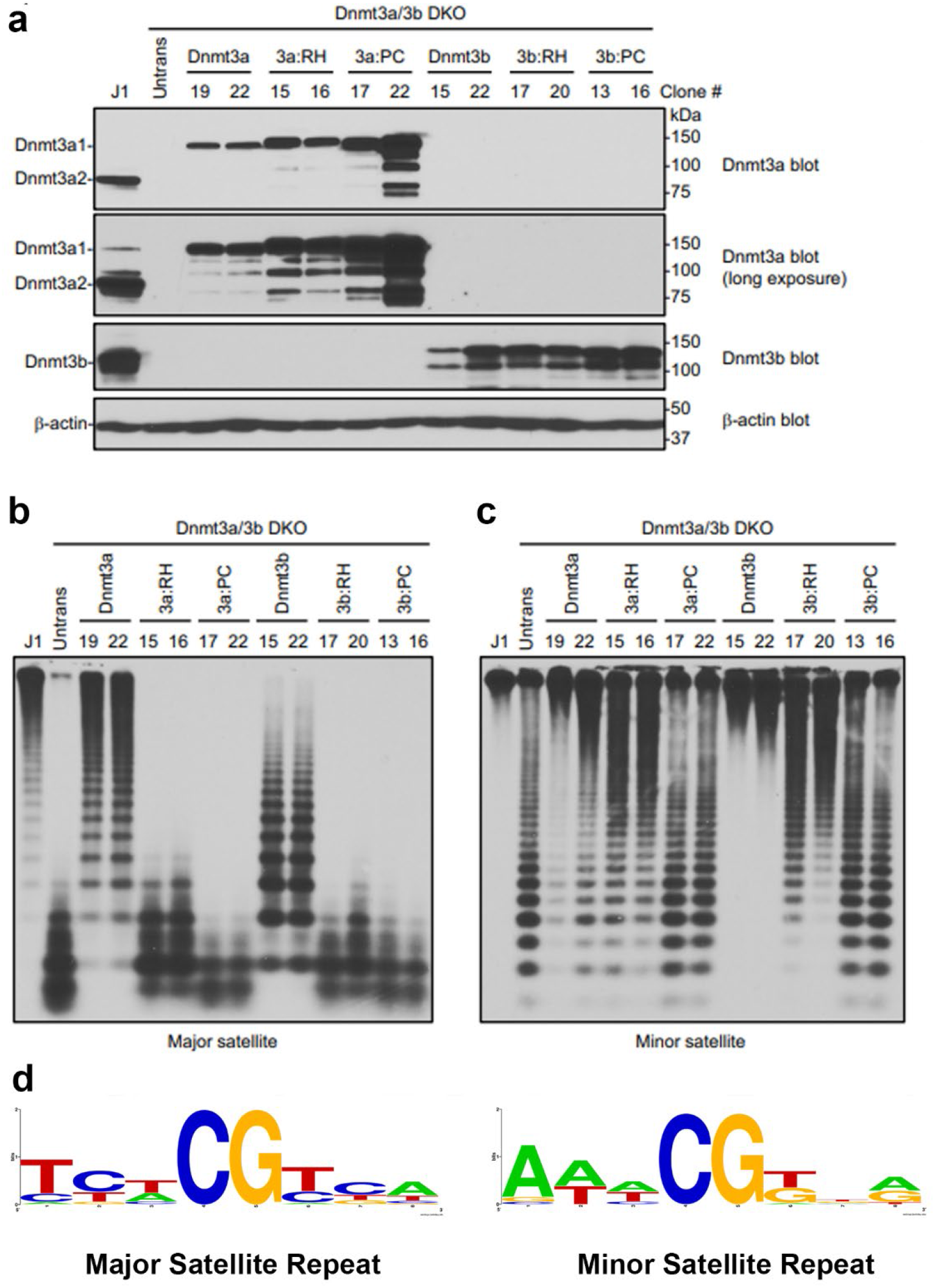
Rescue of DNA methylation at the major and minor satellite repeats in *Dnmt3a/3b* DKO mESCs. **a** *Dnmt3a/3b* DKO mESCs were transfected with plasmids encoding mouse DNMT3A1 WT, DNMT3A1 Arg878His (3A1:RH), DNMT3A Pro705ValCys706Asp (3A1:PC), DNMT3B1 WT, DNMT3B1 Arg829His (3B1:RH), or DNMT3B1 Pro656GlyCys657Thr (3B1:PC), and stable clones were derived. Total cell lysates were used to analyze the expression of DNMT3A or DNMT3B proteins by western blot with anti-DNMT3A, anti-DNMT3B, and anti-β-Actin antibodies. A long exposure of the DNMT3A blot is included to show endogenous DNMT3A1 in WT (J1) mESCs. Note that stable clones showing similar expression levels to those of endogenous DNMT3A or DNMT3B were used for the experiments. **b, c** DNA methylation was analyzed by Southern blot. Genomic DNA was digested with *Mae*II (major satellite repeats) or *Hpa*II (minor satellite repeats), and probed for the major **b** or minor **c** satellite repeats. J1 (WT) and untransfected DKO mESCs were used as controls. The numbers on the top indicate clone#. Complete digestion due to low or no DNA methylation is indicated by low molecular weight bands as seen in untransfected DKOs, and high molecular weight bands as seen in J1 indicate high DNA methylation and protection from digestion. Comparing the activity of DNMT3A1 clones 19/22 with 3A1:RH clones 15/16 at major and minor satellite repeats shows that 15/16 methylate minor repeats similar to 19/22 whereas at major repeats the activity of 15/16 is severely impaired. **d** Consensus sequence of the nucleotides flanking the CpG site in either the major or minor satellite repeats, created using WebLogo shows high prevalence G at N+1 and N+3 positions at minor repeats.

To test if the preference was driven by a potential sequence bias in the major or minor satellite repeats, we analyzed the flanking sequences around CpG sites and computed the consensus sequence logo using WebLogo application^50^ (Fig. 7d). The analysis shows that the major satellite repeats are enriched with CpG sites carrying T at N+1 and A at N+3, and are depleted in CpG sites carrying G at the N+1 and N+3 positions. However, the minor satellite repeats have a high percentage of sites with G at the N+1 and N+3 positions, which are highly preferred by DNMT3B as well as DNMT3A Arg882His. These data confirm that DNMT3A Arg878His acquires catalytic properties similar to DNMT3B, which may allow it to target DNMT3B specific sites in somatic cells and contribute to cancer development.

Our data also show that unlike the WT enzyme, the DNMT3B Arg829His variant was unable to rescue methylation of the major satellite repeats and only partially rescued methylation of the minor satellite repeats (Fig. 7b, c). This is in agreement with our observation that substrates with T at the N+1 position are strongly disfavored by the DNMT3B-C Arg829His variant (Supplementary Fig. 4f). Specifically, about half of the CpG sites in both the major and minor satellite repeats have T at the N+1 position, which could explain the Southern blot results.

Taken together, we propose a model in which substrate specificity and kinetic mechanism together regulate the DNA methylation levels of various genomic regions. At repetitive elements where CpG content is intermediate to high, DNMT3A WT enzyme acts cooperatively to methylate multiple CpGs at its target sites where it prefers A and T at N+3 position. Loss of cooperativity in the DNMT3A WT or DNMT3A Arg882His variant modifies its specificity to be similar to that of DNMT3B where it prefers G at N+3 (Fig. 8). Whereas DNMT3B methylates its targets in a processive manner, DNMT3A WT and Arg882His methylate these sites distributively explaining the lower activity of these enzymes at these sites.

**FIGURE 8.**
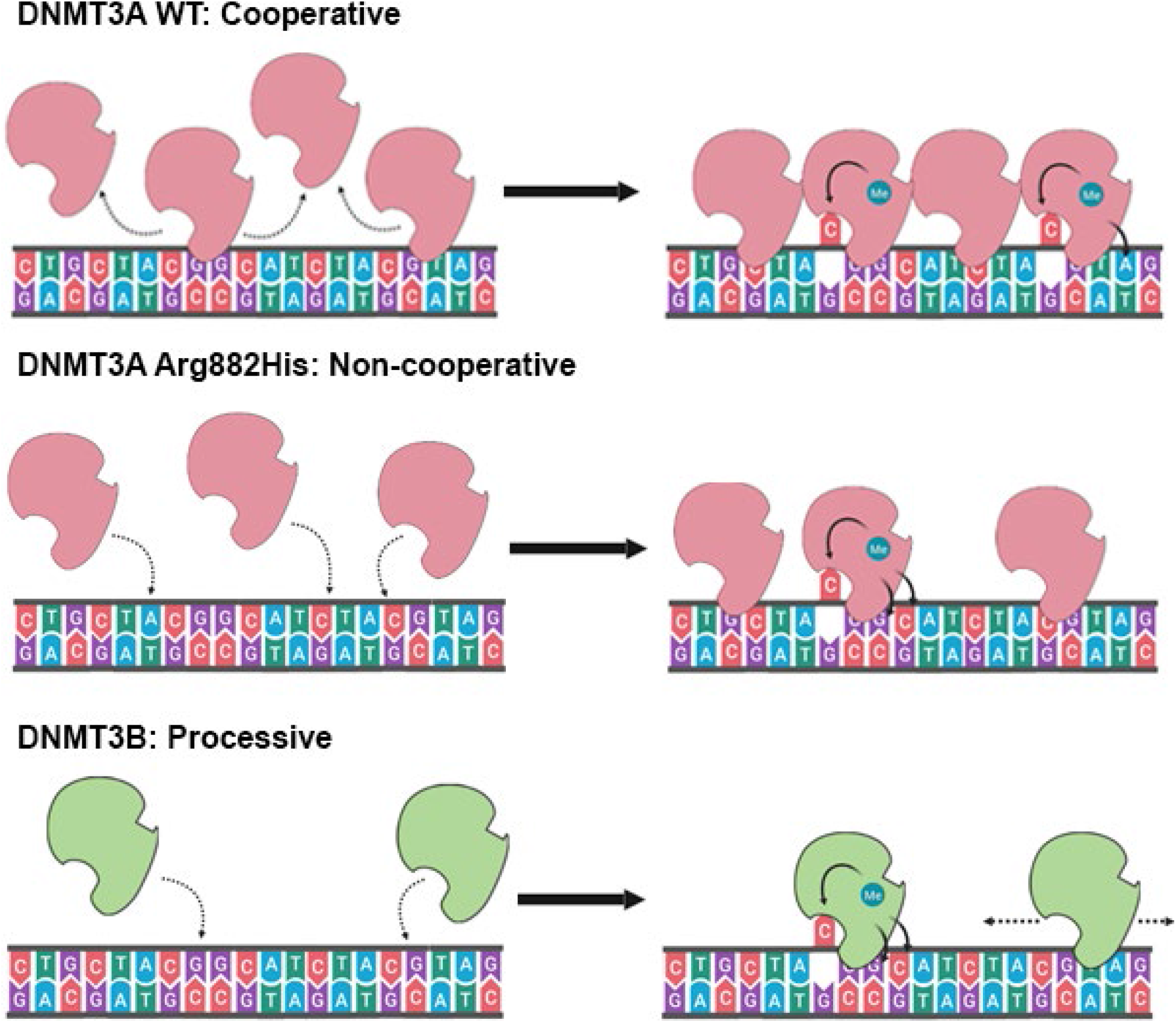
Model showing the different kinetic mechanisms of DNMT3A and DNMT3B and their influence their flanking sequence preferences. At high concentrations, DNMT3A methylates multiple CpG sites rapidly by using cooperative mechanism, and has no strong preference at N+1 and N+3, and a minor preference for A at N+2. However, in absence of cooperative kinetic mechanism at low concentrations of DNMT3A enzyme, for the Arg882His variant, and DNMT3B, the flanking sequence preference for G at the N+1 and C at N+2 position is observed.

## Discussion

Despite numerous studies addressing the biological roles of DNMT3A and DNMT3B in development and diseases, the differences and similarities in their kinetic mechanisms remain poorly understood. Germline mutations in DNMT3A and DNMT3B have deleterious effects and are associated with congenital diseases^17,18^. In AML and other types of leukemia, the majority of somatic mutations in *DNMT3A* affects Arg882, mostly leading to an Arg-to-His substitution^21^. Because of its high prevalence (∼20% in AML) and early occurrence during disease development, the Arg882His variant is considered a founder mutation^53^. Therefore, the variant enzyme has been the subject of many studies. Through these studies, the Arg882His substitution was shown to alter DNA binding properties, attenuate tetramerization, disrupt cooperativity, and change the flanking sequence preference of the DNMT3A enzyme ^22,27,33,40,42^. In this study, we show that disruption of cooperativity alters the flanking sequence preference of DNMT3A-C, suggesting that the gain of flanking sequence preference of the DNMT3A-C Arg882His variant is the consequence of losing the cooperative kinetic mechanism. We systematically computed the temporal order of site preference for the DNMT3A-C, DNMT3A-C Arg882His, DNMT3B-C, and DNMT3B-C Arg829His enzymes. The most important finding in this study is the discovery that the altered flanking sequence preference of the DNMT3A-C Arg882His variant is nearly identical to the preferred substrate sequence of DNMT3B-C. Consequently, we predicted that the DNMT3A Arg882His variant would potentially methylate DNMT3B-specific targets. Our study using *in vitro* bisulfite sequencing and rescue experiments by stable expression of proteins in *Dnmt3a/3b* DKO mESCs cells provide strong evidence supporting this prediction. The data revealed that, although the DNMT3A Arg882His variant has little activity in methylating the major satellite repeats, it’s ability to methylate the minor satellite repeats is almost fully retained. An analysis of the CpG flanking sequence showed an overrepresentation of G at the N+1 and N+3 positions in the minor satellite repeats that is preferred by DNMT3B and the DNMT3A Arg882His variant. The major satellite repeats have high percentages of T and A at the N+1 and N+3 positions, respectively, which are disfavored by DNMT3B WT and the Arg829His variant.

The observations from this study reveal how distinct kinetic properties of DNMT3A and DNMT3B influence the selection of their genomic target sites. Given that under conditions where DNMT3A WT functions as non-cooperative enzyme, its flanking sequence preference is similar to DNMT3A Arg878His variant, we propose that DNMT3A methylates minor satellite repeats using non-cooperative kinetic mechanism. This prediction also explains the rationale behind the lower activity of DNMT3A at minor satellite repeats compared to that of DNMT3B, which methylates its target sites using processive kinetic mechanism. Similarly, although DNMT3A Arg878His variant prefers DNMT3B sites, it methylates these sites using a non-cooperative distributive mechanism explaining an incomplete rescue by the variant enzyme compared to DNMT3B. A high activity of DNMT3A at the major satellite repeats could be justified by the cooperative mechanism used to methylate multiple CpG sites.

These observations also suggest that at genomic regions with sparse or dispersed CpG sites, in the absence of cooperativity, sites with preferred flanking sequence (such as N+1,3=G) could be methylated at higher levels both by DNMT3A and DNMT3B. However, except during early embryogenesis, the tissue specific expression of DNMT3A and DNMT3B ensures that these enzymes methylate distinct regions, besides having many common target sites. The importance of this regulation is highlighted by aberrant expression of DNMT3B in various types of cancer, including AML^14,28–30^. The effect of DNMT3B overexpression in AML is similar to that of DNMT3A Arg882His mutation, leading to an increase in stemness, downregulation in apoptotic genes, and poor patient prognosis^27,29^. Our data here reveal a mechanism by which the DNMT3A Arg882His variant acts like the DNMT3B enzyme, thus providing a mechanistic explanation to above observations in AML patients. We suggest that the oncogenic potential of DNMT3A Arg882His variant may not be only due to its lower activity causing DNA hypomethylation in AML cells, but also due to the gain of DNMT3B-like activity generating aberrant patterns of DNA methylation. Our observations provide novel insights into consequences of cancer-causing mutations on enzymatic activity of DNMT3 enzymes.

## Methods

### Protein Purification

Human DNMT3A-C WT, DNMT3A-C Arg882His, DNMT3A-C Arg882Cys, and DNMT3A-C Arg882Ser, and mouse DNMT3B-C WT and DNMT3B-C Arg829His in pET28a(+) with a 6X His tag, were expressed and purified using affinity chromatography as described^40^. Briefly, transformed BL21 (DE3) pLys cells were induced with 1 mM IPTG at OD_600_ 0.3 and expressed for 2 h at 32°C. Harvested cells were washed with STE buffer (10 mM Tris-HCl (pH 8.0), 0.1 mM EDTA, 0.1 M NaCl), and resuspended in Buffer A (20 mM Potassium Phosphate (pH 7.5), 0.5 M NaCl, 10% (v/v) glycerol, 1 mM EDTA, 0.1 mM DTT, 40 mM imidazole). Cells were disrupted by sonication, followed by removal of cell debris by centrifugation. Clarified lysate was incubated with 0.4 mL Ni-NTA agarose for 3 h at 4°C. The protein bound slurry was packed in a 2 ml Biorad column and washed with 150 ml Buffer A. Protein was eluted using 200 mM imidazole in Buffer A at pH 7.5, then stored in 20 mM HEPES pH 7.5, 40 mM KCl, 1 mM EDTA, 0.2 mM DTT and 20% (v/v) glycerol at −80°C. The purity and integrity of recombinant proteins were checked by SDS-PAGE gel.

### DNA Methylation assays using radiolabeled AdoMet

Radioactive methylation assays to determine kinetic parameters of recombinant enzymes were performed using ^3^H-labelled S-Adenosylmethionine (AdoMet) as a methyl group donor and biotinylated oligonucleotides of varying sizes bound on avidin-coated high-binding Elisa plates (Corning) as described^54^. The DNA methylation reactions were carried out using either 250 nM 30-bp/32-bp DNA substrate in methylation buffer (20 mM HEPES pH 7.5, 50 mM KCl, and 1 mM EDTA, 5 μg/ml BSA). The methylation reaction included 0.76 μM [methyl^3^H] AdoMet (PerkinElmer Life Sciences). Storage buffer was added to compensate for the different enzyme volumes in all reactions. Incorporated radioactivity was quantified by scintillation counting.

### Processivity Assay

Methylation kinetic analyses were performed using two enzyme concentrations and short oligonucleotide 30-bp and 32-bp substrates with 1 and 2 CpG sites, respectively. Low enzyme concentrations relative to DNA substrate concentrations were used to ensure that the reaction occurred under multiple turnover conditions. Each DNA substrate was used at 250 nM, and a 1:1 ratio of labelled and unlabeled AdoMet (final concentration 1.5 μM) was used.

### Cooperativity Assays

To examine cooperativity, increasing concentrations of enzyme were pre-incubated with DNA substrate for 10 minutes at room temperature prior to the addition of AdoMet to start the reaction. AdoMet was a mixture of unlabeled and 0.76 μM ^3^[H] labeled AdoMet to yield a final concentration of 2 μM. Methylation assays were performed using 100 ng of an unmethylated pUC19 plasmid purified from dam^-^ /dcm^-^ *E.coli* strain (C2925I, NEB) or a 1-kb fragment containing 14 CpG sites amplified from the *Meis1* enhancer were used as DNA substrates for filter binding assays. Briefly, 10 μl reaction mix was spotted on 0.5 in DE81 filter that was then washed 5 times in 0.2 M Ammonium Bicarbonate (NH_4_HCO_3_), washed 2-3 times with 100% ethanol, and air dried. Incorporated radioactivity was quantified by scintillation counting^55^.

### Flanking sequence preference

*In vitro* DNA methylation reactions were carried out using 100ng of a 509-bp DNA fragment amplified from the *Suwh1* promoter, or 100 ng of a 1-kb DNA fragment amplified from the *Meis1* enhancer region. Methylation reactions were carried out in methylation buffer (20 mM HEPES pH 7.5, 50 mM KCl, 1 mM EDTA, and 0.05 mg/ml BSA using varying concentrations of each enzyme at 37°C. Samples were taken at 10, 30, or 60 minutes and reaction was stopped by freeze/thaw. DNA methylation was analyzed by bisulfite sequencing as described below.

Flanking sequence preference was also measured using short oligonucleotides and DNA methylation assays using radiolabeled AdoMet as described above. Sixteen different 30-bp substrates were used with varying combinations of the second and third nucleotide around the CpG site on either side (Supplementary Table 3).

### Bisulfite sequencing

Bisulfite conversion was performed using EpiTect Fast Bisulfite Conversion Kit (Qiagen, 59802) according to the manufacturer’s protocol. PCR amplifications were performed with primers as described (Supplementary Fig. 2A-B)^39^. The pooled samples were sequenced using NGS on Wide-Seq platform. The reads were assembled and analyzed by Bismark and Bowtie2. Methylated and unmethylated CpGs for each target were quantified, averaged, and presented as percent CpG methylation.

### Rescue experiments in mESCs

WT (J1) and *Dnmt3a/3b* DKO mESCs were cultured on gelatin-coated petri dishes in mESC medium (DMEM supplemented with 15% fetal bovine serum, 0.1 mM nonessential amino acids, 0.1 mM β-mercaptoethanol, 50 U/mL penicillin, 50 μg/mL streptomycin, and 10^3^ U/mL leukemia inhibitory factor)^9^. For the generation of stable clones expressing DNMT3A or DNMT3B proteins, mESCs were transfected with plasmid vectors using Lipofectamine 2000 (Invitrogen) and then seeded at low density on dishes coated with feeder cells, selected with 6 μg/mL of Blasticidin S HCl (Gibco) for 7-10 days, and individual colonies were picked. Southern blot analysis of DNA methylation at the major and minor satellite repeats were performed as previously described^11,26^.

### Data Analysis

#### Comparative flanking sequence preference analysis

To compare the flanking sequence preference of different enzymes, we first calculated average percent methylation of all CpG sites in the substrate. The fractional variance (v) at each CpG site was calculated by dividing in the percent methylation of each site by average methylation. The preference for a site by an enzyme (B) over enzyme (A) was calculated as relative change (v_B_ − v_A_)/ v_A_, Positive values indicate a preference of enzyme B for a site and values greater than or equal to 1 were considered significant given the preference is more than 2 fold (Supplementary Table 1).

To determine the fractional distribution of nucleotides at preferred sites the occurrence of each nucleotide at N+1/2/3 positions was calculated as a fraction. At each position, the number of times each nucleotide occurred was divided by the total number of preferred sites toc compute the fractional occurrence.

#### Individual flanking sequence preference analysis

Analysis of the bisulfite sequencing data was performed to determine optimal flanking sequence preferred by each enzyme. To take into account the uneven distribution of nucleotides flanking the CpG site, the occurrence of each nucleotide at the N+1/2/3 positions (p1_n_, p2_n_, and p3_n_) was calculated by dividing the number of times it occurred by the total number of CpG sites (s(p1_n_), s(p2_n_), s(p3_n_)). The expected occurrence of a trinucleotide set (O) was computed by multiplying the occurrence of three nucleotides as described by equation (1). This created an expected value, which would predict the probability at which this trinucleotide set would be methylated if there were no flanking sequence preference by the enzyme. With the data obtained from bisulfite sequencing, the fractional methylation (f) for each CpG site was calculated by dividing percent methylation at a site by sum of percent methylation at all sites as described by equation (2). Fractional methylation was sorted by nucleotides at position N+1/2/3 (f(p1_n_), f(p2_n_), and f(p3_n_)) and summed for each nucleotide at a particular position (Σf(p1_n_), Σf(p2_n_), and Σf(p3_n_)). The methylation for the flanking trinucleotide set (M) was calculated then multiplying the summed value for a nucleotide at the three positions as described by equation (3). This gives us the value for observed methylation value at a CpG with a specific flanking trinucleotide. The preference for a flanking trinucleotide by an enzyme was calculated by determining the fold change between the observed and expected values as described by equation (4) (Supplementary Table 2).

*Equations*

v = fractional variance

f = fractional methylation

n = any nucleotide

A,C,G,T = specific nucleotides

s = fraction (number of sites out / 56)

p1 = nucleotide at position N+1

p2 = nucleotide at position N+2

p3 = nucleotide at position N+3

M = observed methylation based on fractional methylation

O = normalized occurrence of each site based on their frequency in the DNA substrate

Σ = sum

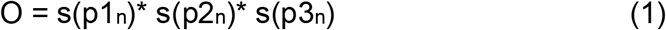

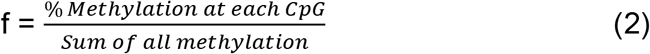

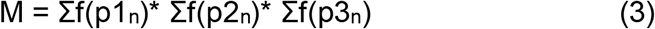

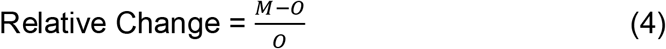

#### Consensus sequence analysis of major and minor satellite repeats

From the mm9 genome, three regions, chr9: 3000466 – 3028144, chr9: 3033472 – 3037264, and chr2: 98506820 – 98507474, were used for the sequences of the major satellite repeats. The minor satellite repeat sequences were obtained from chr2:98505036-98505275 and chr2:98506495-98506615. Additionally, more sequences were obtained from GenBank and used for analysis (accession no. X14462.1, X14463.1, X14464.1, X14465.1, X14466.1, X14468.1, X14469.1, X14470.1). Consensus sequences of the CpG sites and flanking regions in the major and minor satellite repeats were built using WebLogo (https://weblogo.berkeley.edu/logo.cgi).

#### Line and Bar graphs

Data were analyzed using the Prism software (GraphPad). For cooperativity graphs, data were fit with nonlinear, second order polynomial regression curves. Errors were calculated as standard error of the mean (SEM) for two to four independent experiments, as described in the figure legends.

## Acknowledgements

We are thankful to Gowher lab members for discussions. This work was supported by grants from NIH (R01GM118654-01 to H.G. and R01AI12140301A1 to T.C.) and NSF (1716678 to H.G.).

## Author Contributions

A.B.N. and H.G. conceived the project and designed the experiments. A.B.N., L.A., A.R.M., and N.E.F. purified recombinant proteins and performed radiolabeled methylation assays. A.B.N. performed in vitro methylation assays and bisulfite conversion. A.B.N. and H.G. analyzed the data, generated figures, and wrote the manuscript. B.L. performed the rescue experiments in mESCs. T.C. supervised the cellular work and also participated in study design and manuscript writing.

