## Supplemental Table 2 for "The Acute Myeloid Leukemia variant DNMT3A Arg882His is a DNMT3B-like enzyme"

**Trinucleotide preference of DNMT3A-C WT on the 509 bp substrate at 10 minutes**

| Base | N + 1 | N + 2 | N + 3 | DNMT3A WT % | Fractional Methylation (f) |
|---|---|---|---|---|---|
| 71 | G | C | T | 9.5 | 0.009399426 |
| 89 | C | T | C | 34.9 | 0.034530523 |
| 94 | G | C | A | 30.5 | 0.030177105 |
| 100 | C | A | C | 7.1 | 0.007024834 |
| 104 | G | C | C | 21.3 | 0.021074503 |
| 112 | C | G | G | 5.5 | 0.005441773 |
| 114 | G | T | G | 4.8 | 0.004749184 |
| 120 | C | G | A | 19.8 | 0.019590383 |
| 122 | A | C | C | 9.2 | 0.009102602 |
| 127 | C | A | G | 10.5 | 0.010388839 |
| 137 | G | C | T | 24.4 | 0.024141684 |
| 142 | C | C | A | 37.6 | 0.037201939 |
| 176 | C | T | G | 10.6 | 0.010487781 |
| 185 | C | G | C | 13.4 | 0.013258138 |
| 187 | C | C | A | 29.6 | 0.029286633 |
| 192 | C | C | C | 38.5 | 0.038092411 |
| 198 | T | C | C | 45.5 | 0.045018304 |
| 212 | G | A | T | 14.4 | 0.014247551 |
| 228 | C | G | A | 29.9 | 0.029583457 |
| 230 | A | G | C | 12.6 | 0.012466607 |
| 238 | G | C | C | 15.7 | 0.015533788 |
| 244 | C | A | C | 25.1 | 0.024834273 |
| 255 | G | A | G | 5.7 | 0.005639656 |
| 261 | C | T | C | 19.1 | 0.018897794 |
| 267 | G | G | G | 11.7 | 0.011576135 |
| 278 | G | C | G | 8.7 | 0.008607896 |
| 281 | G | C | G | 8.8 | 0.008706837 |
| 284 | A | C | T | 8.7 | 0.008607896 |
| 290 | C | C | T | 52.1 | 0.051548432 |
| 306 | G | C | G | 22.2 | 0.021964975 |
| 309 | C | C | G | 16.4 | 0.016226378 |
| 312 | C | C | G | 18.8 | 0.01860097 |
| 315 | A | C | G | 9.2 | 0.009102602 |
| 318 | C | G | C | 8.3 | 0.00821213 |
| 320 | C | A | C | 14.3 | 0.01414861 |
| 324 | C | C | T | 41.6 | 0.041159592 |
| 334 | C | G | C | 20.1 | 0.019887207 |
| 336 | C | G | C | 18.4 | 0.018205204 |

| | | | | | |
|---|---|---|---|---|---|
| 338 | C | G | T | 14.8 | 0.014643317 |
| 340 | T | C | C | 24.5 | 0.024240625 |
| 357 | A | T | T | 13.5 | 0.013357079 |
| 375 | G | C | G | 15.3 | 0.015138023 |
| 378 | C | G | A | 13.5 | 0.013357079 |
| 380 | A | G | A | 10.5 | 0.010388839 |
| 386 | G | C | G | 6.4 | 0.006332245 |
| 389 | C | G | G | 5.6 | 0.005540714 |
| 391 | G | C | T | 11.3 | 0.01118037 |
| 397 | A | G | C | 30.5 | 0.030177105 |
| 412 | C | G | C | 31.3 | 0.030968636 |
| 414 | C | G | C | 18.4 | 0.018205204 |
| 416 | C | T | C | 25.3 | 0.025032156 |
| 424 | C | C | C | 17.9 | 0.017710498 |
| 436 | A | C | G | 13.9 | 0.013752845 |
| 439 | G | C | T | 12.1 | 0.011971901 |
| 447 | C | G | G | 5.7 | 0.005639656 |
| 449 | G | G | G | 5.7 | 0.005639656 |
| | | | SUM | 1010.7 | |

| Occurrence (O) | No. of sites /56 (s) |
|---|---|
| N + 1 position (p1) | $s(p1_n)$ |
| A ($p1_A$) | 0.142857143 |
| C ($p1_C$) | 0.517857143 |
| G ($p1_G$) | 0.303571429 |
| T ($p1_T$) | 0.035714286 |
| N + 2 position (p2) | $s(p2_n)$ |
| A ($p2_A$) | 0.107142857 |
| C ($p2_C$) | 0.464285714 |
| G ($p2_G$) | 0.321428571 |
| T ($p2_T$) | 0.107142857 |
| N + 3 position (p3) | $s(p3_n)$ |
| A ($p3_A$) | 0.125 |
| C ($p3_C$) | 0.375 |
| G ($p3_G$) | 0.321428571 |
| T ($p3_T$) | 0.178571429 |

| Methylated (M) | | |
|---|---|---|
| N + 1 position (p1) | $\Sigma f(p1_n)$ | |
| A ($p1_A$) | 0.106955575 | $\Sigma f(p1_A)$ |
| C ($p1_C$) | 0.597704561 | $\Sigma f(p1_C)$ |
| G ($p1_G$) | 0.226080934 | $\Sigma f(p1_G)$ |
| T ($p1_T$) | 0.069258929 | $\Sigma f(p1_T)$ |
| N + 2 position (p2) | $\Sigma f(p2_n)$ | |
| A ($p2_A$) | 0.076283764 | $\Sigma f(p2_A)$ |
| C ($p2_C$) | 0.543880479 | $\Sigma f(p2_C)$ |
| G ($p2_G$) | 0.272781241 | $\Sigma f(p2_G)$ |
| T ($p2_T$) | 0.107054517 | $\Sigma f(p2_T)$ |
| N + 3 position (p3) | $\Sigma f(p3_n)$ | |
| A ($p3_A$) | 0.169585436 | $\Sigma f(p3_A)$ |
| C ($p3_C$) | 0.446621154 | $\Sigma f(p3_C)$ |
| G ($p3_G$) | 0.183536163 | $\Sigma f(p3_G)$ |
| T ($p3_T$) | 0.200257247 | $\Sigma f(p3_T)$ |

| Trinucleotide Seq | Occurrence (O= $s(p1_n)*s(p2_n)*s(p3_n)$) |
|---|---|
| AAA | 0.001912625 |
| AAC | 0.005737875 |
| AAG | 0.004911621 |
| AAT | 0.002738879 |
| ACA | 0.008294 |
| ACC | 0.024882 |
| ACG | 0.021298992 |
| ACT | 0.011877008 |
| AGA | 0.005737875 |
| AGC | 0.017213625 |
| AGG | 0.014734863 |
| AGT | 0.008216637 |
| ATA | 0.001912625 |
| ATC | 0.005737875 |
| ATG | 0.004911621 |
| ATT | 0.002738879 |
| CAA | 0.006848 |
| CAC | 0.020544 |
| CAG | 0.017585664 |
| CAT | 0.009806336 |
| CCA | 0.029696 |
| CCC | 0.089088 |
| CCG | 0.076259328 |
| CCT | 0.042524672 |
| CGA | 0.020544 |
| CGC | 0.061632 |
| CGG | 0.052756992 |
| CGT | 0.029419008 |
| CTA | 0.006848 |
| CTC | 0.020544 |
| CTG | 0.017585664 |
| CTT | 0.009806336 |
| GAA | 0.004052625 |
| GAC | 0.012157875 |
| GAG | 0.010407141 |
| GAT | 0.005803359 |
| GCA | 0.017574 |
| GCC | 0.052722 |

| | |
|---|---|
| GCG | 0.045130032 |
| GCT | 0.025165968 |
| GGA | 0.012157875 |
| GGC | 0.036473625 |
| GGG | 0.031221423 |
| GGT | 0.017410077 |
| GTA | 0.004052625 |
| GTC | 0.012157875 |
| GTG | 0.010407141 |
| GTT | 0.005803359 |
| TAA | 0.0004815 |
| TAC | 0.0014445 |
| TAG | 0.001236492 |
| TAT | 0.000689508 |
| TCA | 0.002088 |
| TCC | 0.006264 |
| TCG | 0.005361984 |
| TCT | 0.002990016 |
| TGA | 0.0014445 |
| TGC | 0.0043335 |
| TGG | 0.003709476 |
| TGT | 0.002068524 |
| TTA | 0.0004815 |
| TTC | 0.0014445 |
| TTG | 0.001236492 |
| TTT | 0.000689508 |

| Methylated ( M = $\Sigma f(p1_n)*\Sigma f(p2_n)*\Sigma f(p3_n)$) | Fold Change ((M-O)/O) |
|---|---|
| 0.001727035 | 1.69313967 |
| 0.003251841 | 1.364222573 |
| 0.001937353 | 0.135002855 |
| 0.001913086 | 1.220831198 |
| 0.013750278 | 0.98310352 |
| 0.025890453 | 0.74090418 |
| 0.015424785 | -0.164236381 |
| 0.015231572 | 0.635317401 |
| 0.006666135 | 0.495241151 |
| 0.012551692 | 0.312625156 |
| 0.007477936 | -0.369842198 |
| 0.007384266 | 0.233013732 |
| 0.001870038 | 0.45999212 |
| 0.003521102 | 0.281681141 |
| 0.002097771 | -0.384697629 |
| 0.002071494 | 0.203946488 |
| 0.00617995 | 1.094252843 |
| 0.011636252 | 0.838478672 |
| 0.006932543 | -0.11739336 |
| 0.006845705 | 0.726973948 |
| 0.04920342 | 0.542110954 |
| 0.092645315 | 0.353770682 |
| 0.055195405 | -0.350089282 |
| 0.054504022 | 0.271663759 |
| 0.023853818 | 0.162736959 |
| 0.044914449 | 0.020730189 |
| 0.026758732 | -0.509973514 |
| 0.02642355 | -0.041177647 |
| 0.006691665 | 0.13532643 |
| 0.012599763 | -0.003332652 |
| 0.007506575 | -0.521525469 |
| 0.007412547 | -0.063781064 |
| 0.005580643 | 0.544628536 |
| 0.010507814 | 0.355980788 |
| 0.006260253 | -0.349028267 |
| 0.006181836 | 0.273739821 |
| 0.044431868 | 0.137393029 |
| 0.083660941 | -0.001518449 |

| | |
|---|---|
| 0.049842775 | -0.520654517 |
| 0.049218439 | -0.062076894 |
| 0.02154057 | -0.142416498 |
| 0.040558824 | -0.247154429 |
| 0.024163777 | -0.638578075 |
| 0.023861099 | -0.292814919 |
| 0.006042734 | -0.162633295 |
| 0.011377888 | -0.264902119 |
| 0.006778617 | -0.647098288 |
| 0.006693708 | -0.309486202 |
| 0.000579974 | 0.540693657 |
| 0.001092036 | 0.352526481 |
| 0.000650604 | -0.350686591 |
| 0.000642454 | 0.27049502 |
| 0.004617631 | 0.134495566 |
| 0.008694555 | -0.004062041 |
| 0.005179965 | -0.52187563 |
| 0.00511508 | -0.064466215 |
| 0.002238627 | -0.144601157 |
| 0.00421512 | -0.249072272 |
| 0.002511247 | -0.639498783 |
| 0.002479791 | -0.294616444 |
| 0.000627998 | -0.164766453 |
| 0.001182459 | -0.266774751 |
| 0.000704475 | -0.647997291 |
| 0.000695651 | -0.311245258 |
