## Supplementary Information for "The Acute Myeloid Leukemia variant DNMT3A Arg882His is a DNMT3B-like enzyme"

### SUPPLEMENTARY FIGURES

#### Supplementary Figure 1

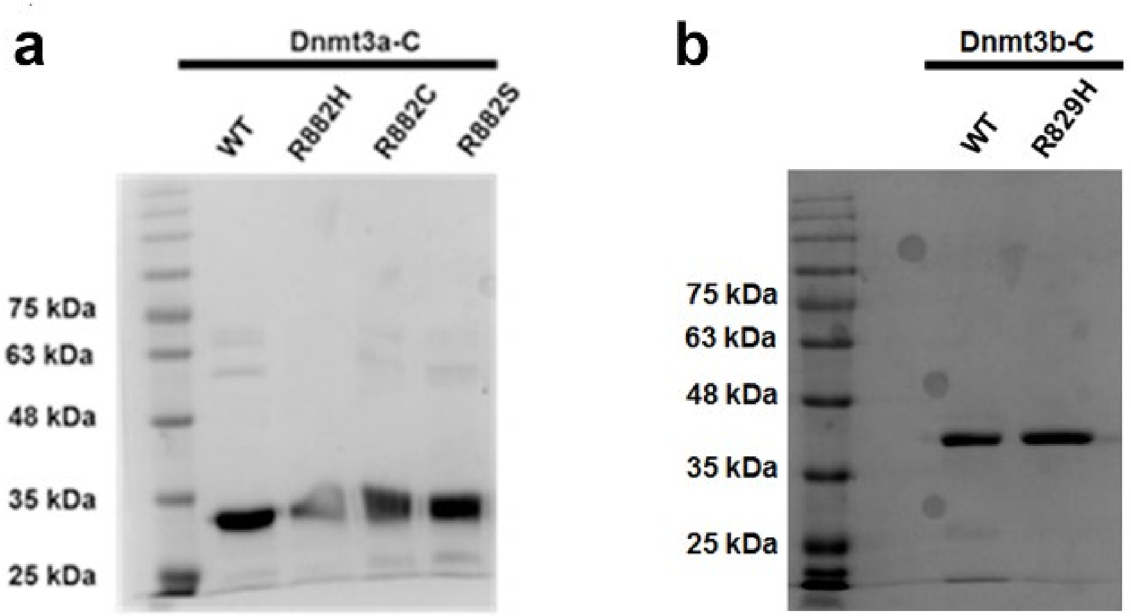


#### Supplementary Figure 2

**509-bp DNA Substrate**

AGATTAGGGAAGGGGGTGTGCGTGAGGGGATCGTGGGACCTGGTTCTGCCCTGCACTCCCCCAC
AGAAAGCGGCTGTATTGCAAAGGCCGCTCCGGCAGCGCACGGCCAAGCGCGGTGTCGCGACCCG
CAGGGCTGCGGCTCGCCATGAAATCCCTCAAGTCTCCCCTCAGGGAACGCTGAAGCCGCGCCAC
GCCCCCGTCCTTACCAGTCCGGATCAGCTGCTGTTCGCGAGCTGCCGGCCACGCACCAGCCCCG
GAGGCGCTCCCGGGGCACAGCCGGCGGCGACTACGCCTCCTCAGGCCCCCGGCGCCGCCGACGC
GCACGCCTCCACACGCGCGCGTCCAGTGGAGACCTGCGATTGGCTGCCAGGTGCCGGCGCGAGA
TCGGCGCGGCTCCGAGCTAGGAGCATGCGCGCGCTCTGACGCCCCTGGTGGCGACGGCTGGACG
CGGGGTTAAATTGAGAAGGAGGAGGGCAGCAGCAATACCCCTGAGGCCTTGAAAGGATCTT

**Supplementary Figure 3**

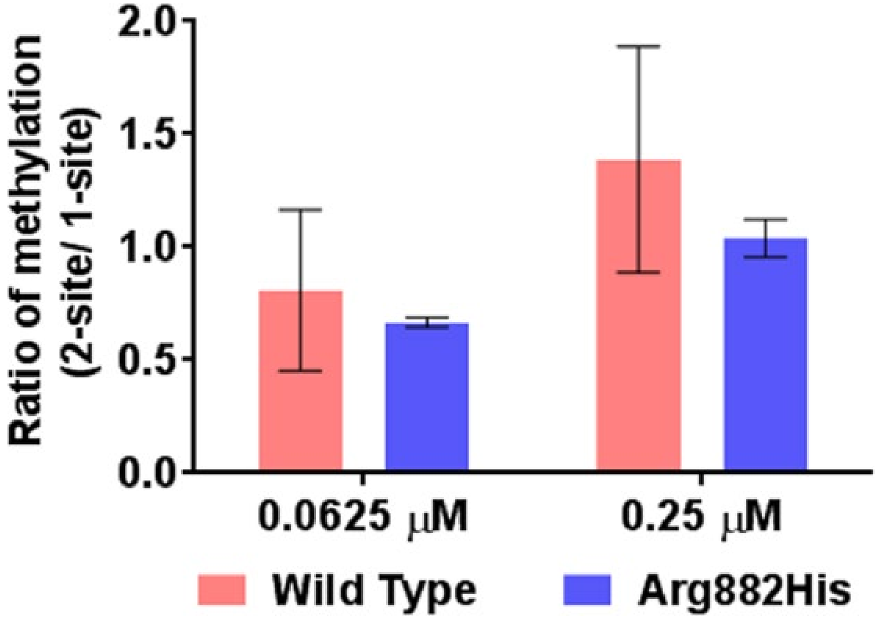

**Supplementary Figure 4**

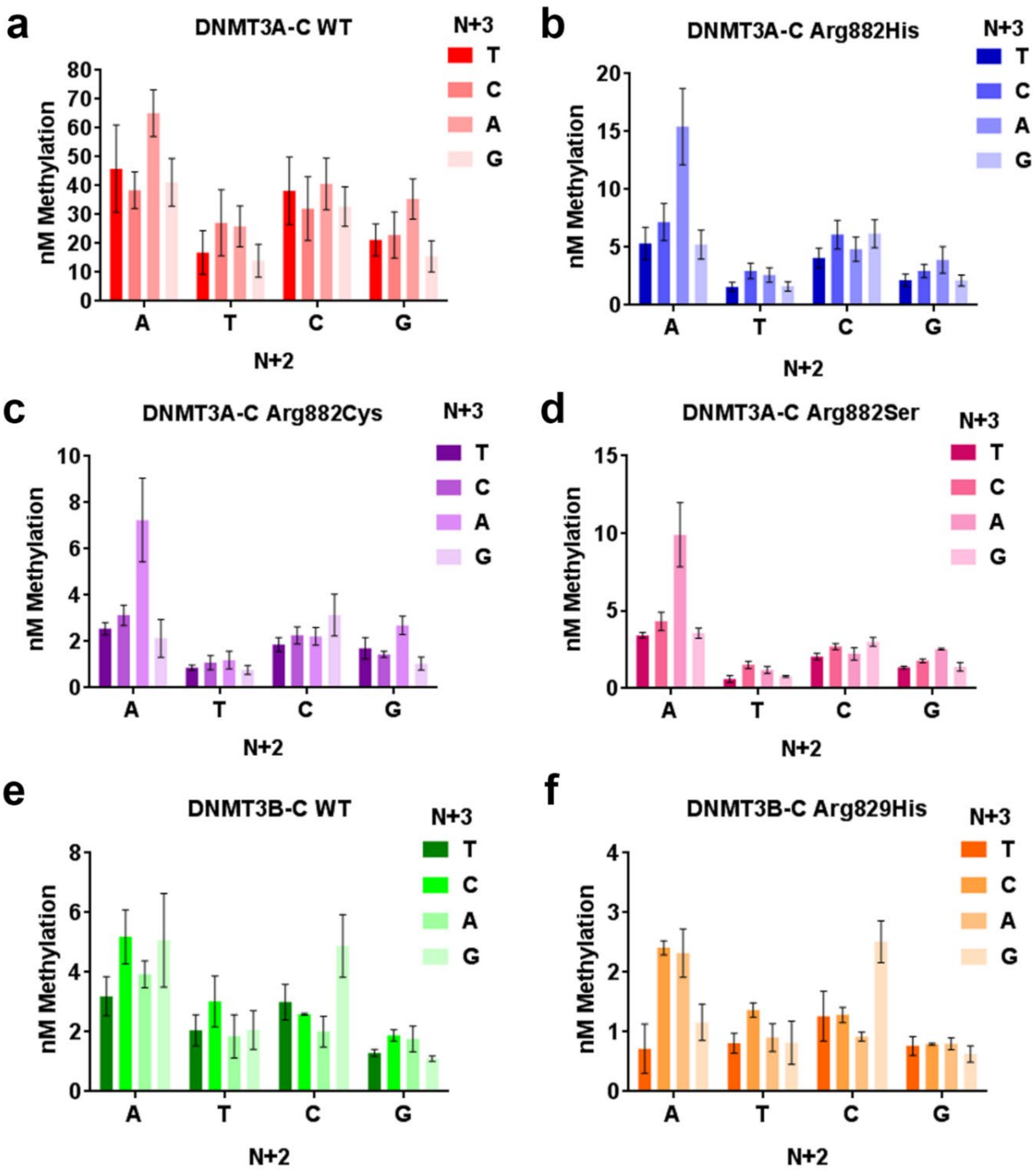

### Supplementary Figure 5

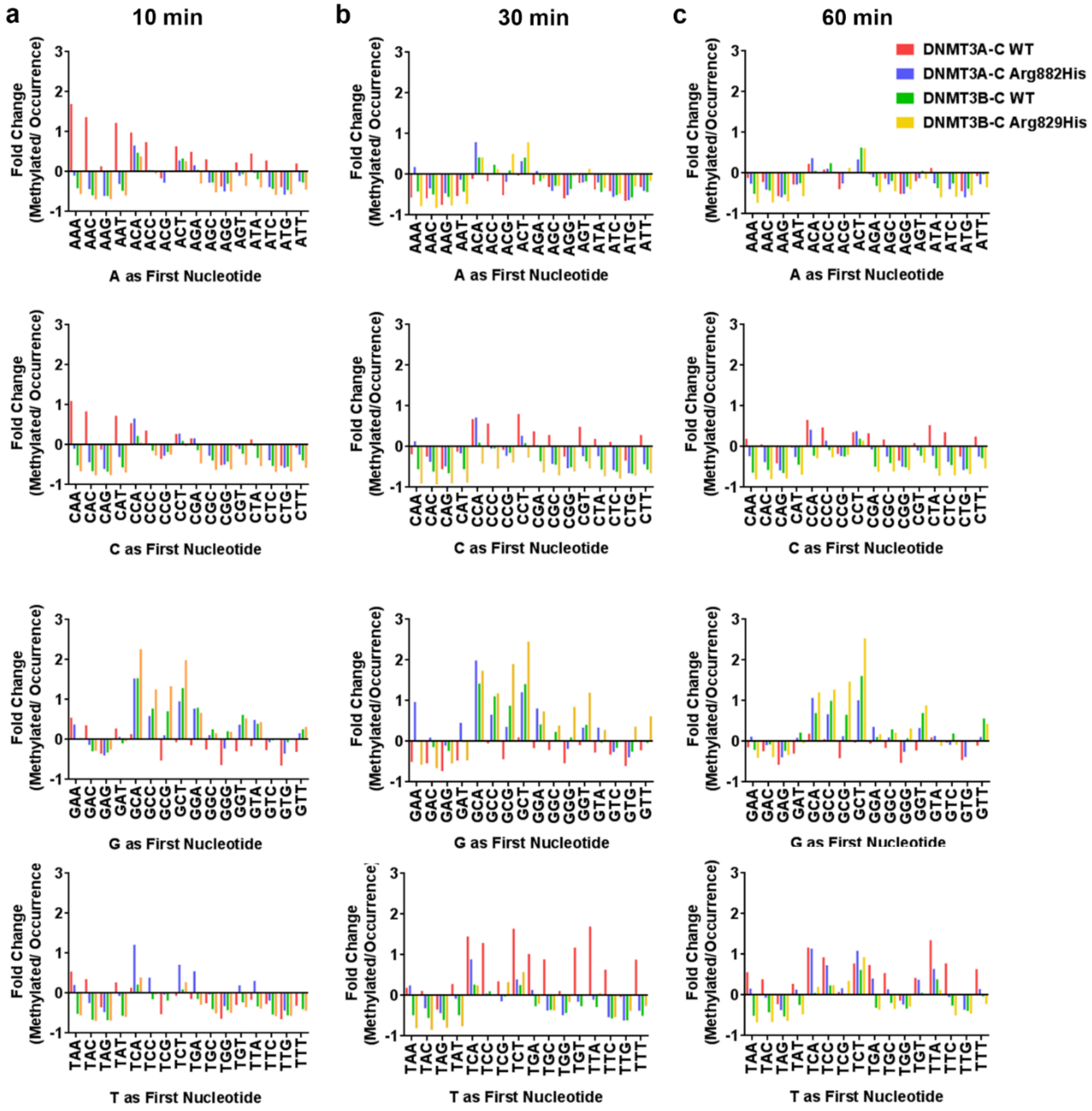

**Supplementary Figure 6**

**a**

### Meis1 DNA Substrate

**GGAAAGCAACGCGCTCAAGCTCAGCTA**GGGTTACATGGTGAGACGCAA
AGCAAAGCAGCCAGAAACAAAGATTAAATCTCTTCTAAATACACATAT
TCCTGGTACTCAGAAACAGCTCTGAATGTTGCTGAAGTTTGGCTCTCC
TGAAATTGGTCATGAGATGTTGGCTTGCTAGGTGAATTTAAGTGTGCT
GAATGGTGACAAGAATGTGTTAGTGGTTAACCACTGTGGACAGACTTC
AGTGTTAGTCTACCCGTTAGCAGTCTCATGACAGCTACAGAAAGTTAA
GAGAAGACAATGATTCAGCACTTGAAGAAAACGCCTCCTGCTCTGATG
TGCTGCCTTCATTTTCCCATTCGAAAGTGATCTCACTGCTGATAAAGC
CGTCATTCCCGACAGCTCTTACTACCATCGCCACATTTAACCAAAGGA
CACTGTGGCCTGAGTACCATGTCCGTGCAGAGCACAAACGGCAAACAA
CACATCTCAGCCACACCACGCTGAACAAGCTCCAGCTCATCCAGGCCT
GGAAGCGAAGCAAAACCGGGCATGGGTCATACTCCAATCGTACACAAG
CTATGGACACCATGAGAAAGAATTTTGGTTTCCATTAAGCATTCCAAT
TGGAGGGCACAGAAGCTAGGCAACGTAATCACAATGATAAATGGAGAG
TTGCAAACTCGTGGACTTAATCACACAGCAATACAATTCCATGAGTGT
CTGAAGTAAGATCTAGCCTAGGCTAGAGGGCACTTGCAGGAGACTCTG
AAGAGCAGGCAAAAGTAAAGGCAGAGGACAGACGGAAGCATTCCAGAC
TATAGGACAGGAAGAGCAGGCCCAGAGGCCTCATGCAGCACAACCCAG
CCATAGAACTCACTGGAACAATAGAGCCAATGATTCTGGACAGTTGGA
GGAACAAGAGCTCTATCCTTGTGGATTGATCCCTATGGTTCTGGGAAT
CCTGACCTGCCAGGGTACA**TCTGTGACGCTGTTAGGGCCAGCAGTT**

**b**

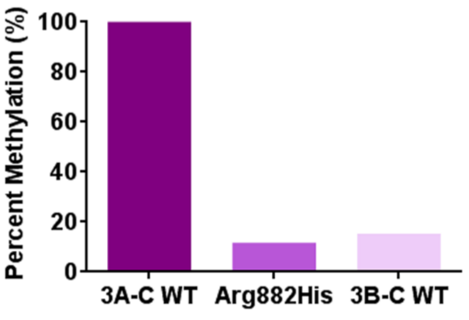


**c**

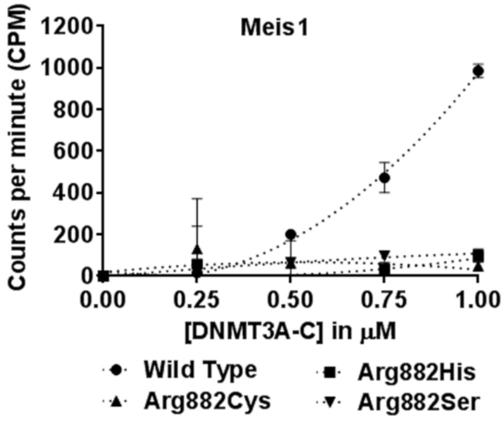

**Tables 1 and 2 are uploaded separately as an Excel file**

**Table 3**

| **30 bp oligomers** |
|---|
| 5' - GAA GAT GGG ATT CAC GTG ACT AGA GTG TAA - 3' |
| 5' - GAA GAT GGG ATA AAC GTT TCT AGA GTG TAA - 3' |
| 5' - GAA GAT GGG ATA CAC GTG TCT AGA GTG TAA - 3' |
| 5' - GAA GAT GGG ATT TAC GTA ACT AGA GTG TAA - 3' |
| 5' - GAA GAT GGG ATT GAC GTC ACT AGA GTG TAA - 3' |
| 5' - GAA GAT GGG ATG CAC GTG CCT AGA GTG TAA - 3' |
| 5' - GAA GAT GGG ATC TAC GTA GCT AGA GTG TAA - 3' |
| 5' - GAA GAT GGG ATG AAC GTT CCT AGA GTG TAA - 3' |
| 5' - GAA GAT GGG ATG TAC GTA CCT AGA GTG TAA - 3' |
| 5' - GAA GAT GGG ATG GAC GTC CCT AGA GTG TAA - 3' |
| 5' - GAA GAT GGG ATT AAC GTT ACT AGA GTG TAA - 3' |
| 5' - GAA GAT GGG ATC AAC GTT GCT AGA GTG TAA - 3' |
| 5' - GAA GAT GGG ATC GAC GTC GCT AGA GTG TAA - 3' |
| 5' - GAA GAT GGG ATA GAC GTC TCT AGA GTG TAA - 3' |
| 5' - GAA GAT GGG ATA TAC GTA TCT AGA GTG TAA - 3' |
| 5' - GAA GAT GGG ATC CAC GTG GCT AGA GTG TAA - 3' |

### SUPPLEMENTARY FIGURE LEGENDS

**Supplementary Figure 1**

Coomassie-stained SDS-PAGE gel showing purified recombinant His-tagged DNMT3 enzymes. **a** DNMT3A-C WT, Arg882His, Arg882C, and Arg882S. **b** DNMT3B-C WT and DNMT3B-C Arg829His.

**Supplementary Figure 2**

Sequence of the 509-bp DNA substrate used for DNA methylation assays. Underlined sequence indicates primer-binding site for bisulfite sequencing.

**Supplementary Figure 3**

Methylation of a 1-site and 2-site substrate at two different enzyme concentrations. All reactions were carried out using 0.25 µM of DNA substrate and a 1:1 ratio of radioactively labelled and unlabeled AdoMet. Samples were incubated for 60 minutes and the incorporation of radioactivity was measured as counts per minute using scintillation counter. The ratio of incorporated methylation of the 2-site substrate compared to the 1-site substrate was plotted at different concentrations for each enzyme.

**Supplementary Figure 4**

Methylation activity using short DNA substrates with varying N +2 and N +3 nucleotide sequences with T at the N +1 position. Each enzyme at 1 µM concentration was incubated with 250 nM 30-bp substrate and a mixture of labeled and unlabeled SAM for

10 minutes. Total methylation activity was plotted for each dinucleotide set grouped by second nucleotide. Error bars represent SEM of three independent experiments with two different enzyme purifications.

**Supplementary Figure 5**

Analysis of the bisulfite data to determine the best trinucleotide set based on the fractional methylation of the 509-bp substrate at 10 **a**, 30 **b**, or 60 minutes **c**. Data are separated based on the N+1 position, and are sorted alphabetically from left to right in each graph.

**Supplementary Figure 6**

**a** Sequence of the 1-kb DNA substrate amplified from the *Meis1* enhancer used for in vitro DNA methylation assays and analysis by bisulfite sequencing. Underlined sequence indicates primer-binding site for bisulfite sequencing. The CpG site in red is the site preferred by DNMT3A-C Arg882His and DNMT3B-C WT compared to DNMT3A-C WT from Fig. 6 a, b. **b** Methylation activity of DNMT3A-C WT and Arg882 variants was measured for 10 minutes with the *Meis1* substrate in the presence of 0.25 to 1 µM enzyme, respectively. The enzymes were pre-incubated with DNA for 10 minutes at room temperature and the reaction was initiated by addition of AdoMet. Total methylation activity was plotted against enzyme concentration using an average and standard deviation ($n \geq 3$ independent experiments). **c** Methylation activity of DNMT3A-C WT, DNMT3A-C Arg882His, and DNMT3B-C WT on the *Meis1* substrate at 10 minutes, measured by bisulfite analysis. The activity of DNMT3A-C WT was set to 100% and the relative activity of DNMT3A-C Arg882His and DNMT3B-C WT was compared to this level.

**Table 1**

Data analysis for Fig. 2b and 3a. To determine fold methylation, the percent methylation at each CpG sites was divided by the average methylation percentage of all CpG sites for each enzyme. Preference of each site by the low concentration DNMT3A-C WT compared to high concentration was determined by calculating the relative change. To determine the fractional distribution of nucleotides at the N+1/2/3 positions, the sequence of the preferred sites was analyzed and fractional occurrence of each nucleotide at each position was calculated and represented in parts of a whole plot in the lower panel Fig. 2a.

**Table 2**

Data for Fig. 5a, DNMT3A-C WT at 10 minutes on the 509-bp substrate. **a** The nucleotide number and flanking sequence preference are listed, then with the percent methylation at each site and the fractional methylation calculated as described in methods by equations 2. **b** The fraction of sites occurring in the substrate are listed at the top by equation 1, and the calculated fractional methylation for each nucleotide at a given position is on the bottom by equation 3. **c** The occurrence (left) and the methylation level of each nucleotide set (middle). The fold change was calculated with equation 4 (right).

**Table 3**

Table containing the sequences of each 30-bp substrate used to determine substrate specificity at the N+2 and N+3 nucleotide positions. To make sure each side of the substrate is methylated equally, the flanking sequencing is the same on each side of the CpG site.
