## Supplemental Table 1 for "The Acute Myeloid Leukemia variant DNMT3A Arg882His is a DNMT3B-like enzyme"

| Base | Site No. | DNMT3A-C WT 1 µM% | Fractional Variance (A) | DNMT3A-C WT 0.25 µM% |
|---|---|---|---|---|
| 71 | 1 | 9.5 | 0.526367864 | 1.5 |
| 89 | 2 | 34.9 | 1.93370931 | 2.4 |
| 94 | 3 | 30.5 | 1.689917879 | 2.6 |
| 100 | 4 | 7.1 | 0.393390719 | 1.6 |
| 104 | 5 | 21.3 | 1.180172158 | 2.3 |
| 112 | 6 | 5.5 | 0.30473929 | 1.1 |
| 114 | 7 | 4.8 | 0.265954289 | 1.1 |
| 120 | 8 | 19.8 | 1.097061443 | 1.3 |
| 122 | 9 | 9.2 | 0.509745721 | 1 |
| 127 | 10 | 10.5 | 0.581775007 | 1.2 |
| 137 | 11 | 24.4 | 1.351934303 | 1.7 |
| 142 | 12 | 37.6 | 2.083308598 | 2.2 |
| 176 | 13 | 10.6 | 0.587315722 | 1.1 |
| 185 | 14 | 13.4 | 0.742455724 | 1.5 |
| 187 | 15 | 29.6 | 1.640051449 | 1.6 |
| 192 | 16 | 38.5 | 2.133175027 | 2.1 |
| 198 | 17 | 45.5 | 2.521025032 | 2.3 |
| 212 | 18 | 14.4 | 0.797862867 | 1.2 |
| 228 | 19 | 29.9 | 1.656673593 | 1.8 |
| 230 | 20 | 12.6 | 0.698130009 | 1.3 |
| 238 | 21 | 15.7 | 0.869892154 | 1.4 |
| 244 | 22 | 25.1 | 1.390719303 | 1.4 |
| 255 | 23 | 5.7 | 0.315820718 | 0.7 |
| 261 | 24 | 19.1 | 1.058276442 | 1.3 |
| 267 | 25 | 11.7 | 0.64826358 | 1.3 |
| 278 | 26 | 8.7 | 0.482042149 | 1.5 |
| 281 | 27 | 8.8 | 0.487582863 | 1.1 |
| 284 | 28 | 8.7 | 0.482042149 | 0.8 |
| 290 | 29 | 52.1 | 2.88671218 | 3.6 |
| 306 | 30 | 22.2 | 1.230038587 | 1.5 |
| 309 | 31 | 16.4 | 0.908677154 | 1.4 |
| 312 | 32 | 18.8 | 1.041654299 | 1.7 |
| 315 | 33 | 9.2 | 0.509745721 | 0.9 |
| 318 | 34 | 8.3 | 0.459879292 | 1.1 |
| 320 | 35 | 14.3 | 0.792322153 | 1.2 |
| 324 | 36 | 41.6 | 2.304937172 | 2.9 |
| 334 | 37 | 20.1 | 1.113683586 | 1.7 |
| 336 | 38 | 18.4 | 1.019491442 | 1.6 |
| 338 | 39 | 14.8 | 0.820025725 | 1.2 |
| 340 | 40 | 24.5 | 1.357475017 | 1.5 |
| 357 | 41 | 13.5 | 0.747996438 | 1.6 |
| 375 | 42 | 15.3 | 0.847729297 | 1.4 |
| 378 | 43 | 13.5 | 0.747996438 | 1.6 |
| 380 | 44 | 10.5 | 0.581775007 | 1.4 |
| 386 | 45 | 6.4 | 0.354605719 | 0.6 |
| 389 | 46 | 5.6 | 0.310280004 | 0.8 |

| | | | | |
|---|---|---|---|---|
| 391 | 47 | 11.3 | 0.626100722 | 0.9 |
| 397 | 48 | 30.5 | 1.689917879 | 1.7 |
| 412 | 49 | 31.3 | 1.734243594 | 2.6 |
| 414 | 50 | 18.4 | 1.019491442 | 1.4 |
| 416 | 51 | 25.3 | 1.401800732 | 1.5 |
| 424 | 52 | 17.9 | 0.99178787 | 1.2 |
| 436 | 53 | 13.9 | 0.770159296 | 0.7 |
| 439 | 54 | 12.1 | 0.670426437 | 0.9 |
| 447 | 55 | 5.7 | 0.315820718 | 0.6 |
| 449 | 56 | 5.7 | 0.315820718 | 0.8 |
| AVG | | 18.04821429 | | 1.471428571 |

| Fractional Variance (B) | Relative Change ((B-A)/A) | Fold Change 2^((B-A)/A) | Sites prefer |
|---|---|---|---|
| 1.019417476 | 0.936701584 | 1.914146941 | |
| 1.631067961 | -0.156508193 | 0.897193956 | |
| 1.766990291 | 0.045607194 | 1.03211748 | |
| 1.087378641 | 1.764118693 | 3.396664423 | |
| 1.563106796 | 0.32447354 | 1.252207407 | |
| 0.747572816 | 1.45315534 | 2.738062427 | |
| 0.747572816 | 1.81090716 | 3.508628402 | Fractional c |
| 0.883495146 | -0.194671227 | 0.873771996 | |
| 0.67961165 | 0.333236598 | 1.259836572 | **A** |
| 0.815533981 | 0.401803051 | 1.321158036 | **C** |
| 1.155339806 | -0.145417197 | 0.904117888 | **G** |
| 1.495145631 | -0.282321576 | 0.822266765 | **T** |
| 0.747572816 | 0.27286362 | 1.208203625 | |
| 1.019417476 | 0.373034705 | 1.295074152 | p-values fo |
| 1.087378641 | -0.336985043 | 0.791694073 | |
| 1.427184466 | -0.330957635 | 0.795008596 | **A** |
| 1.563106796 | -0.379971727 | 0.76845265 | **C** |
| 0.815533981 | 0.022148058 | 1.015470309 | **G** |
| 1.223300971 | -0.261592038 | 0.834166894 | **T** |
| 0.883495146 | 0.265516644 | 1.202066444 | |
| 0.951456311 | 0.093763527 | 1.067150407 | |
| 0.951456311 | -0.315853092 | 0.803375799 | |
| 0.475728155 | 0.506323454 | 1.420425784 | |
| 0.883495146 | -0.16515656 | 0.891831734 | |
| 0.883495146 | 0.362864078 | 1.285976321 | |
| 1.019417476 | 1.114789086 | 2.165633464 | |
| 0.747572816 | 0.533222087 | 1.447157643 | |
| 0.54368932 | 0.127887513 | 1.092692538 | |
| 2.446601942 | -0.152460727 | 0.899714559 | |
| 1.019417476 | -0.171231304 | 0.8880844 | |
| 0.951456311 | 0.047078499 | 1.033170602 | |
| 1.155339806 | 0.109139382 | 1.078584631 | |
| 0.611650485 | 0.199912938 | 1.148629037 | |
| 0.747572816 | 0.625584864 | 1.542836159 | |
| 0.815533981 | 0.029295947 | 1.020513981 | |
| 1.970873786 | -0.144933836 | 0.904420855 | |
| 1.155339806 | 0.037403999 | 1.026265491 | |
| 1.087378641 | 0.066589278 | 1.047237946 | |
| 0.815533981 | -0.005477565 | 0.99621044 | |
| 1.019417476 | -0.24903408 | 0.841459605 | |
| 1.087378641 | 0.453721683 | 1.369568744 | |
| 0.951456311 | 0.122358652 | 1.088513011 | |
| 1.087378641 | 0.453721683 | 1.369568744 | |
| 0.951456311 | 0.635436893 | 1.553408099 | |
| 0.40776699 | 0.149916566 | 1.109505305 | |
| 0.54368932 | 0.752253814 | 1.684422222 | |

| | | |
|---|---|---|
| 0.611650485 | -0.023079732 | 0.984129632 |
| 1.155339806 | -0.316333758 | 0.803108182 |
| 1.766990291 | 0.01888241 | 1.013174315 |
| 0.951456311 | -0.066734382 | 0.954796788 |
| 1.019417476 | -0.272780038 | 0.827723006 |
| 0.815533981 | -0.177713294 | 0.884103211 |
| 0.475728155 | -0.382299015 | 0.767214018 |
| 0.611650485 | -0.087669502 | 0.94104166 |
| 0.40776699 | 0.291134389 | 1.223602016 |
| 0.54368932 | 0.721512519 | 1.648909843 |

rred by DNMT3B-C WT compared to DNMT3A-C WT

| N + 1 | N + 2 | N + 3 |
|---|---|---|
| G | T | G |
| C | A | C |
| C | G | G |
| G | C | G |

istribution of bases at the preferred sites

| N + 1 | N + 2 | N + 3 |
|---|---|---|
| 0 | 0.25 | 0 |
| 0.5 | 0.25 | 0.25 |
| 0.5 | 0.25 | 0.75 |
| 0 | 0.25 | 0 |

r each base at the preferred sites

| N + 1 | N + 2 | N + 3 |
|---|---|---|
